# Cyclin-dependent kinase 5 (Cdk5) activity is modulated by light and gates rapid phase shifts of the circadian clock

**DOI:** 10.1101/2024.02.08.579499

**Authors:** Andrea Brenna, Micaela Borsa, Gabriella Saro, Jürgen A. Ripperger, Dominique A. Glauser, Zhihong Yang, Antoine Adamantidis, Urs Albrecht

## Abstract

The circadian clock enables organisms to synchronize biochemical and physiological processes over a 24-hour period. Natural changes in lighting conditions, as well as artificial disruptions like jet lag or shift work, can advance or delay the clock phase to align physiology with the environment. Within the suprachiasmatic nucleus (SCN) of the hypothalamus, circadian timekeeping and resetting rely on both membrane depolarization and intracellular second-messenger signaling. Voltage-gated calcium channels (VGCCs) facilitate calcium influx in both processes, activating intracellular signaling pathways that trigger *Period* (*Per*) gene expression. However, the precise mechanism by which these processes are concertedly gated remains unknown.

Our study demonstrates that cyclin-dependent kinase 5 (Cdk5) activity is modulated by light and regulates phase shifts of the circadian clock. We observed that knocking down Cdk5 in the SCN of mice affects phase delays but not phase advances. This is linked to uncontrolled calcium influx into SCN neurons and an unregulated protein kinase A (PKA) – calcium/calmodulin-dependent kinase (CaMK) – cAMP response element-binding protein (CREB) signaling pathway. Consequently, genes such as *Per1* are not induced by light in the SCN of Cdk5 knock-down mice. Our experiments identified Cdk5 as a crucial light-modulated kinase that influences rapid clock phase adaptation. This finding elucidates how light responsiveness and clock phase coordination adapt activity onset to seasonal changes, jet lag, and shift work.

---

The circadian system coordinates biochemical and physiological functions in our body, synchronizing them with the environmental day-night cycle. Misalignment of the internal body clock with the external light-dark cycle, caused by shift work or jet lag, leads to inefficient regulation of body functions. Consequently, circadian misalignment can result in obesity, cancer, addictive behaviors, cardiovascular disease, and neurological disorders ^1^. Therefore, it is crucial to understand how the environment impacts the clock and how these entities interact.

In mammals, the master circadian clock is located in the ventral part of the hypothalamus, just above the optic chiasm, in the suprachiasmatic nuclei (SCN). These nuclei coordinate daily cycles of physiology and behavior ^2^. Molecular daily oscillations are generated at the cellular level by a cell-autonomous transcription-translation feedback loop (TTFL) involving a set of clock genes ^3^ and post-translational modifiers such as kinases ^4, 5^. Circuit-level interactions among SCN cells produce a coherent daily oscillation ^6^, which can be modulated by light signals to match the environmental light-dark cycle. Light is perceived by melanopsin-containing intrinsically photosensitive retinal ganglion cells (ipRGCs) in the eye, and the signal produced in these cells travels via the retinohypothalamic tract (RHT) to the SCN ^7^. The release of glutamate at the RHT terminals stimulates AMPA/NMDA receptors, leading to Ca^2+^ influx into the SCN ^8^. Additionally, the activity of various kinases is altered, including DARPP-32 (dopamine and cAMP-regulated phosphoprotein of 32 kD), PKA (protein kinase A), and CaMK (Ca^2+^/calmodulin-dependent kinases). This cascade culminates in the phosphorylation of CREB (cyclic AMP response element binding protein) ^9 - 13^. This event promotes chromatin phosphorylation ^14^ and acetylation ^15^ via the recruitment of CRTC1 (cAMP-regulated transcriptional co-activator 1) and the histone acetyltransferase CBP (CREB-binding protein), involving the clock protein PER2. Consequently, immediate-early gene and clock gene expression is induced ^16, 17, 18^, causing a phase shift of the TTFL in oscillating cells of the SCN ^6^. This manifests at the behavioral level as a change of locomotor activity onset (phase shift) the day after the light pulse. The direction of the phase shift depends on the clock’s temporal state. Light perceived in the early night promotes phase delays, while a light pulse late at night promotes phase advances. Light in the middle of the day does not alter the clock phase ^19^. For this so-called resetting of the circadian clock, the *Per1* and *Per2* genes appear to be important in mice. While *Per1* is essential for phase advances, *Per2* function is necessary for phase delays ^20, 21^.

Voltage-gated calcium channels (VGCCs) are classified into high voltage-activated channels, which include L-type, and low voltage-activated subtypes, also known as T-type channels ^22^. T-type VGCCs are involved in phase delays, whereas L-type VGCCs are related to phase advances ^23, 24^. Ca_V_3.1, Ca_V_3.2, and Ca_V_3.3 belong to the T-type channel family, which is critically important for neuronal excitability ^25^. The activity of these T-type channels is regulated by various kinases, including PKA ^26^, PKC ^27^, and Cdk5 (cyclin-dependent kinase 5) ^28^.

Cdk5 is a proline-directed serine/threonine kinase that forms a complex with its neural activators p35 or p39 ^29, 30^ and cyclin I ^31^. The complex of Cdk5 and its activators controls various neuronal processes such as neurogenesis, neuronal migration, and synaptogenesis ^32,33^. *In vivo* and *in vitro* experiments show that Cdk5 kinase activity is low in the light phase and high in the dark phase ^34, 35^. It regulates the circadian clock in the SCN via phosphorylation of PER2 at serine 394. Upon phosphorylation by Cdk5, PER2 is stabilized and enters the nucleus to participate in the regulation of the TTFL and CREB-related transcriptional events ^15, 34^. Since Cdk5 regulates the T-type channel Ca_V_3.1 ^28^ and the circadian clock via PER2 phosphorylation ^34^, we analyzed the potential role of Cdk5 in the light-mediated clock resetting mechanism.

## Results

### Cdk5 knock-down in the SCN impairs light-induced phase delays

Light perceived during the dark period elicits changes in the clock phase ^19^. To test whether *Cdk5* plays a role in this process, we knocked down *Cdk5* in the SCN via stereotaxic application of adeno-associated viruses (AAVs). We injected an adenovirus expressing shRNA to silence *Cdk5* (shCdk5) and, as a control, an adenovirus expressing a control shRNA (scr) into the SCN ^34^. Consistent with our previous observations ^34^, we found that silencing *Cdk5* in the SCN reduced its expression in the SCN (Supplementary Fig. 1a) and the expression of PER2 (Supplementary Fig. 1b). Under constant darkness (DD) conditions, this knock-down of *Cdk5* shortened the clock period in male mice, as assessed by wheel-running activity (Fig. 1a, b and Supplementary Fig. 1c). This period was not influenced by light pulses (Supplementary Fig. 1d). However, the onset of activity was affected after releasing mice into constant darkness (DD). Light at zeitgeber time (ZT) 14 (where ZT0 is lights on and ZT12 is lights off) delayed the clock phase, whereas light at ZT22 advanced it in control (scr) animals, with light at ZT10 having no effect (Fig. 1a, c, Aschoff type II protocol). The animals with silenced *Cdk5* in the SCN (shCdk5) behaved similarly to controls (scr), except for ZT14. Light did not elicit a phase delay at this time, suggesting that *Cdk5* plays a role in the phase delay mechanism. Similar results were obtained for female animals (Supplementary Fig. 1e-g).

**Figure 1.**
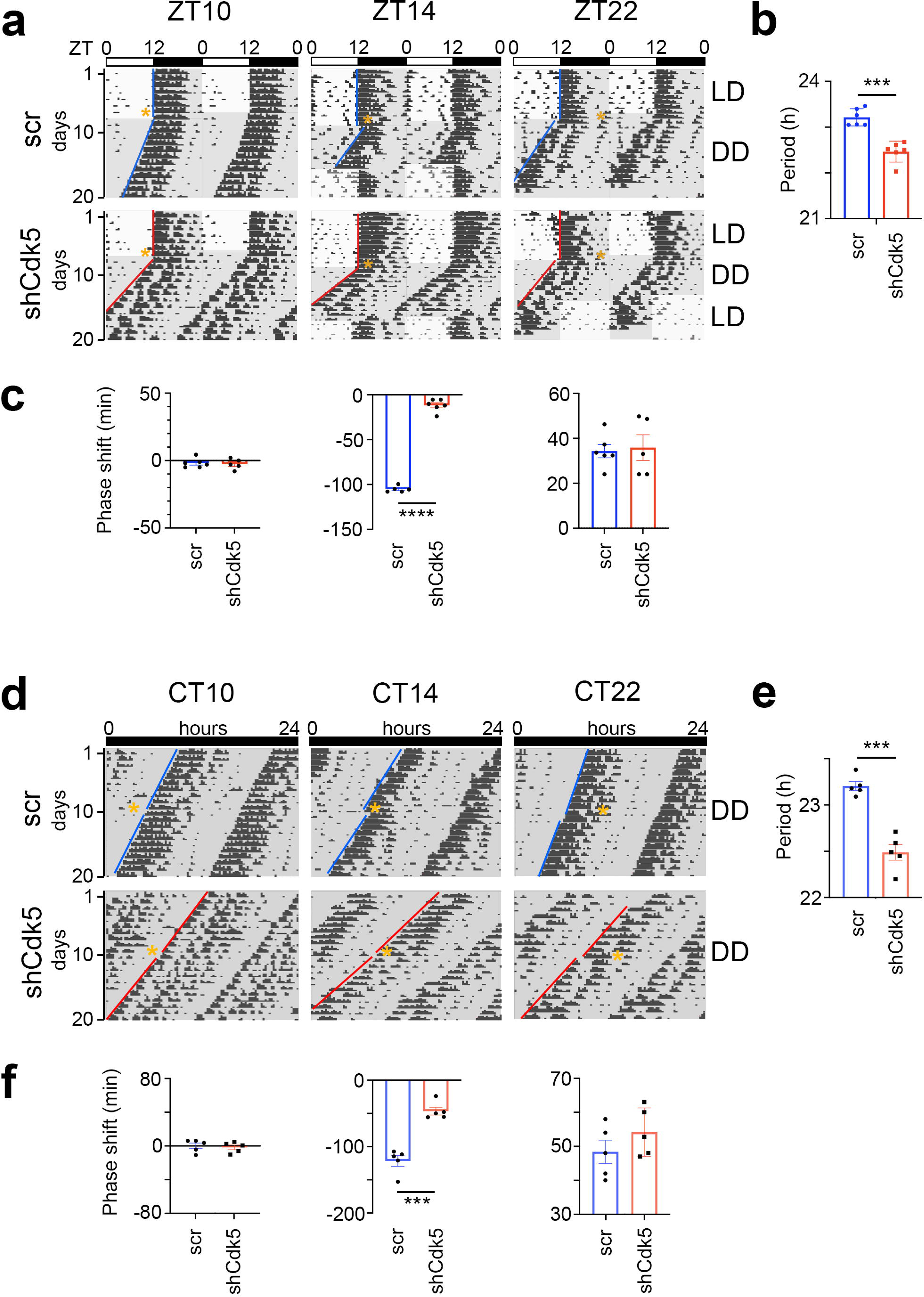
Knock-down of Cdk5 in the SCN shortens period and reduces phase delays but not phase advances. (**a**) Examples of double plotted wheel-running actograms of control (scr) and Cdk5 knock-down (shCdk5) male mice. Animals were kept under a 12 h light / 12 h dark cycle (white and grey areas, respectively) (LD). After 8-10 days they received a 15 min. light pulse at the indicated zeitgeber times (ZT) (yellow stars). After the light pulse animals were released into constant darkness (DD). This light pulse assessment is termed Aschoff type II. (**b**) Circadian period (τ) of shCdk5 mice (red) is significantly shorter compared to scr controls (blue). τ scr = 23.21 ± 0.08 h, τ shCdk5 = 22.47 ± 0.09 h. All values are mean ± SEM, unpaired t-test with Welch’s correction, n = 6, ***p < 0.001. (**c**) Quantification of phase shifts (φ) after a 15 min. light pulse at ZT10, ZT14 and ZT22. The phase shift at ZT14 is strongly reduced in shCdk5 animals (red) compared to scr controls (blue). scr: φ ZT10: -1.93 ± 1.43 min., φ ZT14: -105.24 ± 1.54 min., φ ZT22: 34.30 ± 2.97 min., shCdk5: φ ZT10: -2.60 ± 1.72 min., φ ZT14: -11.80 ± 2.81 min., φ ZT22: 35.88 ± 5.68 min. All values are mean ± SEM, unpaired t-test with Welch’s correction, n = 5-6, ****p < 0.0001. (**d**) Examples of double plotted wheel-running actograms of control (scr) and Cdk5 knock-down (shCdk5) mice. Animals were kept under DD. After 8-10 days they received a 15 min light pulse at the indicated circadian times (CT) (orange stars). This light pulse assessment is termed Aschoff type I. (**e**) Circadian period of shCdk5 mice (red) is significantly shorter compared to scr controls (blue). τ scr = 23.20 ± 0.05 h, τ shCdk5 = 22.48 ± 0.09 h. All values are mean ± SEM, unpaired t-test with Welch’s correction, n = 5, ***p < 0.001. (**f**) Quantification of phase shifts after a 15 min. light pulse at CT10, CT14 and CT22. The phase shift at CT14 is strongly reduced in shCdk5 animals (red) compared to scr controls (blue). scr: φ ZT10: 0.12 ± 3.31 min., φ ZT14: -121.52 ± 8.18 min., φ ZT22: 48.40 ± 3.43 min., shCdk5: φ ZT10: -1.68 ± 2.78 min., φ ZT14: -46.60 ± 5.84 min., φ ZT22: 54.16 ± 3.19 min. All values are mean ± SEM, unpaired t-test with Welch’s correction, n = 5, ***p < 0.001.

To corroborate our observations, we performed the same experiment in DD (Aschoff type I protocol). The shCdk5 animals displayed a shorter period compared to scr controls (Fig. 1d, e), consistent with previous observations ^34^. After determining each animal’s clock period, we administered light pulses of 15 minutes at circadian times (CT) 10, CT14, and CT22 for each animal (orange stars, Fig. 1d). Light at CT10 had no effect on both the shCdk5 and scr control mice (Fig. 1f). Light applied at CT14 promoted a phase delay in scr control mice.

However, silencing of *Cdk5* impaired the delay of the clock phase (Fig. 1d, f), which is consistent with the observation at ZT14 (Fig. 1a, c). Light at CT22 elicited normal phase advances in shCdk5 and scr controls (Fig. 1d, f), similar to the light pulse applied at ZT22 (Fig. 1a, c). From these experiments, we conclude that *Cdk5* plays a role in delaying the clock phase in response to a light pulse in the early activity period of mice.

### Cdk5 activity is modulated by light in the early night

Given that Cdk5 plays a significant role in the phase shift of the circadian clock, we investigated whether the light signal at ZT14 could affect the levels of Cdk5 and its co-activator p35 in the SCN. To this end, we collected SCN samples at ZT14 in the dark or after a 15-minute light pulse and performed a western blot on total protein extracts. To ensure proper light induction, we measured the light-dependent phosphorylation of PKA (Fig. 2a, b), CaMKII and CREB (Supplementary Fig. 2a, b, c, d). We confirmed that PKA, CaMKII and CREB phosphorylation levels increased in response to light in the SCN (Fig. 2a, b, Supplementary Fig. 2a, b, c, d). Interestingly, we observed that light could also increase the p35 protein level, although the levels of Cdk5 remained unaffected (Fig. 2a, c). Given the increase in p35 levels due to light, we wondered whether this event would affect the kinase activity of the Cdk5/p35 complex. We performed an *in vitro* kinase assay using immunoprecipitated Cdk5 from SCN tissue collected from mice either not exposed to light or exposed to light at ZT14. We used the recombinant histone H1 as a substrate in the presence of radioactive ATP ^34^. Surprisingly, our results indicated that Cdk5 kinase activity decreased in response to light (Fig. 2d, e), suggesting that light may affect the interaction between Cdk5 and p35. To test this hypothesis, we performed a co-immunoprecipitation experiment using an antibody against Cdk5. Our results revealed that SCN extracts from mice that received a light pulse at ZT14 contained less p35 in a complex with Cdk5 (Fig. 2f).

**Figure 2.**
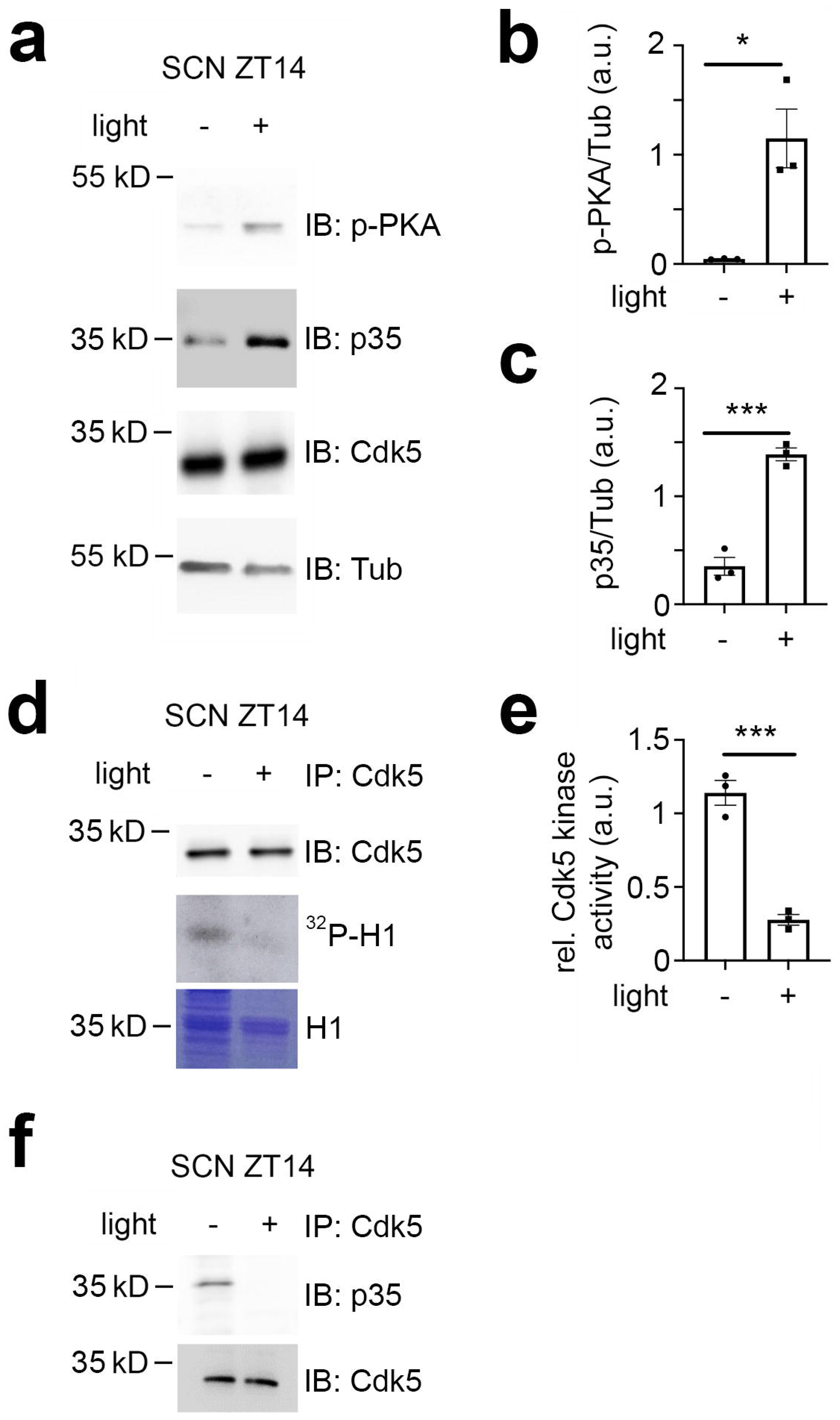
Cdk5 activity is modulated by light in the early night. Immuno Western blotting (IB), immunoprecipitation (IP) and Cdk5 kinase activity assays from SCN tissue extracts harvested 30 min after light (+) and no light (-) given at ZT14. (**a**) Western blot depicting the amounts of phospho PKA (p-PKA), p35 co-activator, Cdk5, and tubulin (Tub, control) before and after light pulse at ZT14. (**b**) Quantification of p-PKA relative to tubulin. Values are the mean ± SEM. Unpaired t-test, n = 3, *p < 0.05. (**c**) Quantification of p35 co-activator of Cdk5. Values are the mean ± SEM. Unpaired t-test, n = 3, ***p < 0.001. (**d**) Cdk5 kinase activity assay. IP of SCN extracts with antibodies against Cdk5 showing the presence of Cdk5 (upper panel) and total protein with Coomassie blue staining (lower panel) as a control for the presence of H1. The middle panel depicts histone H1 phosphorylated by Cdk5, visualized as ^32^P-histone H1 (^32^P-H1). (**e**) Quantification of Cdk5 kinase activity relative to H1 levels. Values are the mean ± SEM. Unpaired t-test, n = 3, ***p < 0.001. (**f**) Co-immunoprecipitation of p35 with Cdk5 before and after a light pulse.

Taken together, the results support the hypothesis that light affects Cdk5 activity by interfering with the formation of the Cdk5/p35 complex. Interestingly, the light pulse at ZT14 might affect more than just Cdk5/p35 protein-protein interactions, potentially involving additional unknown proteins (Supplementary Fig. 2e).

### Cdk5 impacts the CREB signaling pathway via calcium/calmodulin-dependent kinases (CaMK)

Deletion of a cAMP-responsive element (CRE) in the *Per1* promoter blunted light-induced *Per1* expression in the SCN at night ^36^. Because nocturnal light induces phosphorylation of CRE binding protein (CREB) and phosphorylated CREB (p-CREB) can bind to CREs ^37, 38, 39^, we investigated whether Cdk5 is involved in the pathway evoking the CREB phosphorylation at serine-133 (pSer-133), a site known to be involved in phase delays, and *Per1* induction ^11^. Therefore, we performed immunohistochemical analysis using an antibody detecting phosphate on CREB at serine 133 (p-CREB-S133) (Fig. 3a, Supplementary Fig. 3b, control Supplementary Fig. 3c). In the SCN of control animals (scr), we observed p-CREB-S133 in nuclei of neurons after the light was delivered at ZT14 but not in controls (Fig. 3a, arrowheads). In contrast, p-CREB-S133 was already detected in nuclei before the light pulse in shCdk5 animals (Fig. 3a, arrowheads), indicating that *Cdk5* plays a role in gating the phosphorylation of CREB.

**Figure 3.**
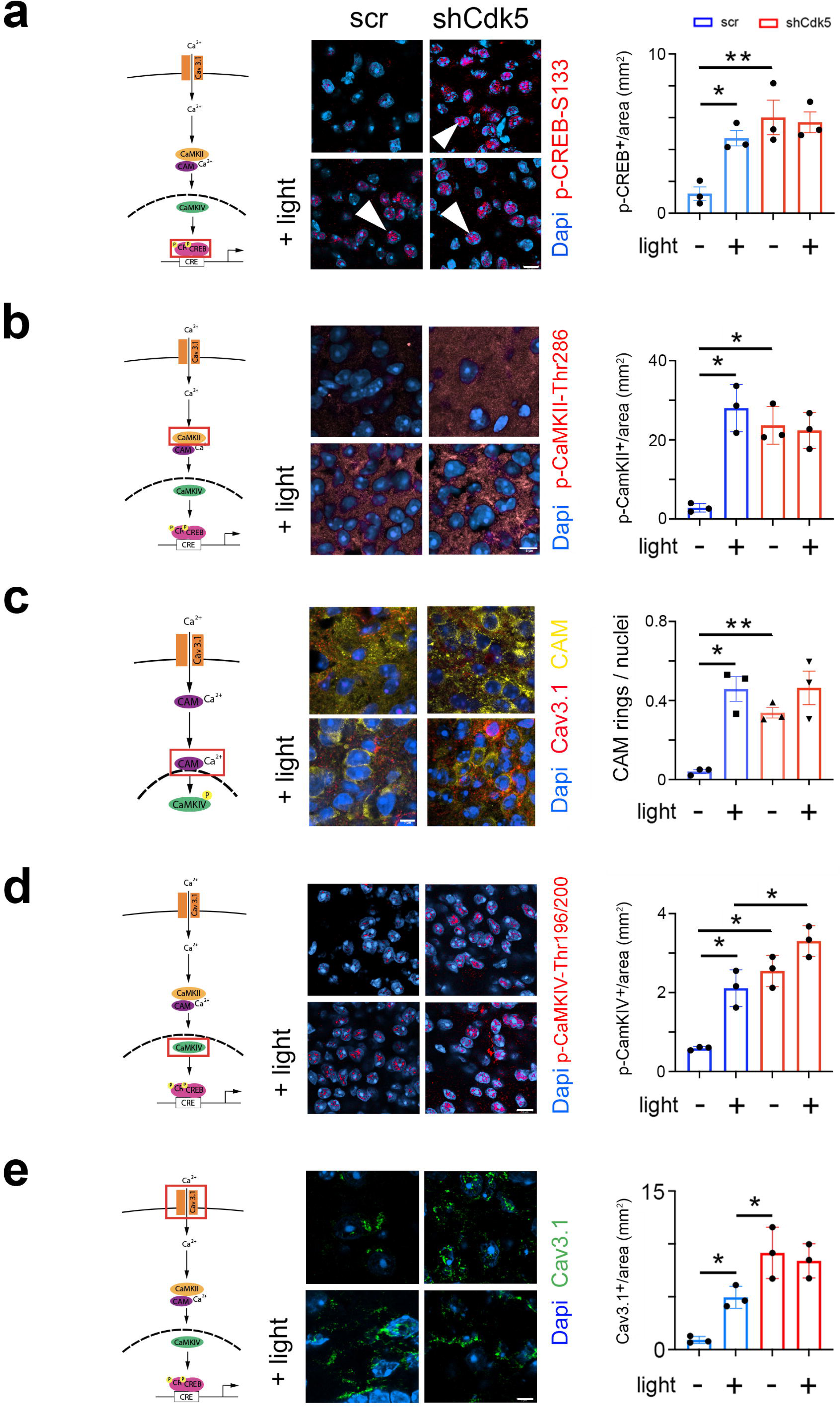
Cdk5 impacts on the CREB signaling pathway via calcium/calmodulin dependent kinases (CaMK). The cartoons on the left of each figure depict the CaMK pathway with the red rectangle indicating the visualization of a particular the component. (**a**) Immunohistochemistry on the SCN of control (scr) and shCdk5 mice using an antibody recognizing phospho-serine 133 of CREB (p-CREB-S133) before and after a light pulse at ZT14. The red color shows p-CREB-S133 and the blue color represents Dapi stained nuclei of SCN cells. Scale bar: 8 µm. The right panel shows the quantification of the p-CREB-S133 signal. Values are the mean ± SEM. Unpaired t-test with Welch’s correction, n = 3, *p < 0.05. (**b**) Immunohistochemistry on the SCN of control (scr) and shCdk5 mice using an antibody recognizing Cam kinase II (CaMKII) before and after a light pulse at ZT14. The red color shows CaMKII and the blue color represents Dapi stained nuclei of SCN cells. Scale bar: 8 µm. The right panel shows the quantification of the CaMKII signal. Values are the mean ± SEM. Unpaired t-test with Welch’s correction, n = 3, *p < 0.05. (**c**) Translocation of calmodulin (CAM) in response to a light pulse at ZT14 in SCN neurons of control (scr) and Cdk5 knock-down (shCdk5) animals. CAM (yellow) accumulates around the nuclei in scr controls. In shCdk5 SCN neurons, this accumulation around the nuclei is already seen before the light pulse which is clearly different from scr controls. Scale bar: 7 µm The right panel shows the quantification of CAM rings. Values are the mean ± SEM. Unpaired t-test with Welch’s correction, n = 3, *p < 0.05, **p < 0.01. (**d**) Immunohistochemistry on the SCN of control (scr) and shCdk5 mice using an antibody recognizing Cam kinase IV (CaMKIV) before and after a light pulse at ZT14. The red color shows CaMKIV and the blue color represents Dapi stained nuclei of SCN cells. Scale bar: 5 µm. The right panel shows the quantification of the CaMKIV signal. Values are the mean ± SEM. Unpaired t-test with Welch’s correction, n = 3, *p < 0.05. (**e**) Immunohistochemistry on the SCN of control (scr) and shCdk5 mice using an antibody recognizing the calcium channel Cav3.1 before and after a light pulse at ZT14. The green color shows Cav3.1 and the blue color represents Dapi stained nuclei of SCN cells. Scale bar: 5 µm. The right panel shows the relative Cav3.1 signal. Values are the mean ± SEM. Unpaired t-test with Welch’s correction, n = 3, *p < 0.05.

The CREB/CRE transcriptional pathway has been shown to be activated by calcium/calmodulin-dependent kinase II (CaMKII) and mitogen-activated protein kinase (MAPK) ^40, 41, 42^. Pharmacological inhibition of CaMKII but not of MAPK affected light-induced phase delays in hamsters ^43^. Therefore, we tested whether phosphorylated CaMKII (p-CaMKII) is affected by the knock-down of *Cdk5* in the SCN of mice. We observed that p-CaMKII presence (alpha isoform) in the cytoplasm of SCN cells increased after light at ZT14 compared to no light in control animals (Fig. 3b, left panels). In shCdk5 SCN, however, p-CaMKII was already present before the light pulse in significantly higher levels than controls (Fig. 3b, control Supplementary Fig. 3d). This result indicates that *Cdk5* is gating the phosphorylation of CaMKII alpha.

CaMKII has been shown to shuttle Ca^2+^/calmodulin (Ca^2+^/CAM) to the nucleus to trigger CREB phosphorylation and gene expression ^44^. Therefore, we investigated whether CAM localization was influenced by a light pulse and whether *Cdk5* plays a role in this process. We observed that in control animals, CAM was distributed evenly in the cytoplasm of cells in SCN tissue before a light pulse. However, after the light pulse, it was localized around the nuclei (Fig. 3c). Interestingly, in the SCN of shCdk5 animals, CAM was already localized around the nuclei before the light administration and remained there after the light pulse, suggesting that *Cdk5* is gating CAM localization in the cell.

Once delivered to the nucleus, Ca^2+^/CAM triggers a highly cooperative activation of the nuclear CaMK cascade, including CaMKIV, to rapidly phosphorylate CREB for the transcription of target genes ^44, 45^. Therefore, we tested whether a light pulse affected the phosphorylation of CaMKIV and whether this was influenced by *Cdk5*. In control animals, we detected p-CaMKIV to be strongly present in the SCN after, but not before, a light pulse (Fig. 3d, control Supplementary Fig. 3e). In shCdk5 SCN, p-CaMKIV was always detectable, independent of the light pulse (Fig. 3d). This indicated that Cdk5 was gating phosphorylation of CaMKIV.

Calcium entry is regulated by channels, such as T-type VGCC, which are involved in phase delays ^23^. Previous reports show that Cdk5 directly or indirectly can phosphorylate Cav3.1 *in vitro* ^28^. Thus, we looked at the influence of light and *Cdk5* on the T-type channel Cav3.1 using immunohistochemical staining. We observed that the level of Cav3.1 protein was significantly increased on the surface of SCN cells after the light pulse (Fig. 3e, blue bars). This suggests that light inhibits internalization and degradation of this channel. Interestingly, in the *Cdk5*-depleted SCN cells, Cav3.1 staining was already high on the cell surface before the light signal (Fig. 3e, red bars). We observed no difference in the Cav3.1 signal between SCN samples obtained from shCdk5 mice before and after the light pulse (Fig. 3e, red bars), suggesting that *Cdk5* may be directly or indirectly involved in the regulation of Cav3.1 localization. This is consistent with previously described effects of Cdk5 on the cellular localization of other receptors such as the D2 and TRPV1 receptors ^46, 47^.

### Cdk5 modulates neuronal activity in response to light at ZT14

Neuronal activity in response to light at ZT14 requires calcium influx. At night, neuronal cell membranes are hyperpolarized, creating a Ca^2+^ gradient. A light stimulus at night promotes membrane depolarization and VGCC activation, which evokes a Ca^2+^ influx into SCN neurons, ultimately changing the phase of the circadian clock ^48, 49^. Our results shown in figure 3 indicate that Cdk5 regulates the gating between light and the CaMKII pathway, which relies on Ca^2+^ availability. Thus, we tested whether Cdk5 regulated the light-mediated Ca^2+^ influx into SCN neurons. To this end, we employed *in vivo* calcium imaging to assess changes in calcium levels in the SCN in freely moving mice after 15 minutes of a light pulse given at ZT14. First, we injected an adeno-associated virus (AAV) expressing the shCdk5 sequence into the SCN to silence *Cdk5*. This AAV co-expresses the calcium indicator GCaMP7 under the neuron-specific synapsin 1 promoter. As a control, we injected an AAV carrying a non-specific shRNA (scrambled sequence) instead of shCdk5 (see Materials and Methods). Consistent with our previous results, the construct expressing shCdk5 in the SCN produced a shortened free-running period in mice (Supplementary Fig. 4a-c). Animals injected with AAV were implanted with a chronic optical fiber placed above the SCN to allow for longitudinal imaging of GCaMP7 signals using fiber photometry. After habituation, ΔF/F_0_ (or the ratio of change in GCaMP7 fluorescence to the baseline fluorescence, see methods) was recorded before and after light pulse delivery at ZT14 in both groups of mice (Fig. 4a).

**Figure 4.**
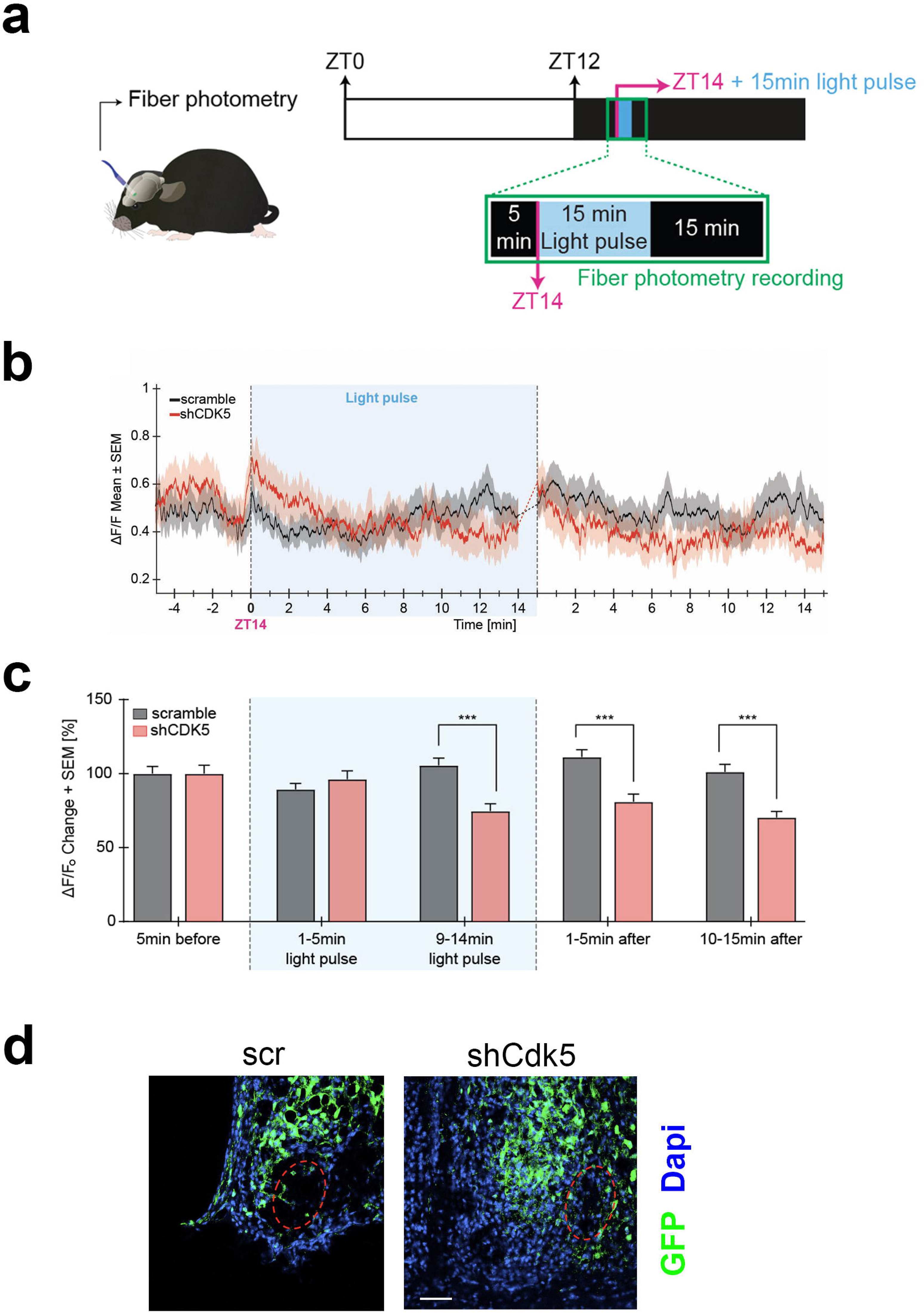
Neuronal activity in response to light at ZT14 is modulated by CDK5. (**a**) Illustration of the chronic optic fiber implantation in the SCN for fiber photometry recording in freely moving mice (left). The animals were previously infected either with AAV9-hSyn1-chl[1x(shNS)]-jGCaMP7b-WPRE-SV40p(A)(scr) or AAV9-hSyn1-chl[mouse(shCdk5)]-jGCaMP7b-WPRE-SV40p(A)(shCdk5). The experimental timeline of one trial is shown on the right. White and dark boxes represent the light and dark phase, respectively. Fiber photometry recordings were done in the 5 min. before the light pulse, during the light pulse, and 15 min. after. The light pulse was 15 min. long and was delivered at ZT14 (blue box; dashed lines between minutes 14-15 are not included in the analysis). (**b**) Mean traces ± SEM of cell activity (normalized ΔF/F_0_) of GCaMP7b-expressing SCN neurons (black: scr, red: shCdk5) 5 min. before, 15 min. during, and 15 min. after the light pulse (± 20 s) light pulse delivered at ZT14. N = 15 trials, n = 5 mice / red = shCdk5, N = 12 trials, n = 4 mice. (**c**) Bar plot showing the percentage of ΔF/F_0_ changes ± SEM 5 min. before the light pulse, in the first and last 5 min. during the light pulse and the first and last 5 min. after the light pulse. *9-14 minutes of light pulse*: scramble (black bar) 105.5 ± 19.3 ΔF/F_0_ vs. shCdk5 (red bar) 74.7 ± 17.2 ΔF/F_0_. 1*-5 minutes after light pulse*: scramble (black bar) 111.2 ± 19.3 ΔF/F_0_ vs. shCdk5 (red bar) 81.0 ± 17.8 ΔF/F_0_. Black = scr, N = 15 trials, n = 5 mice / red = shCdk5, N = 12 trials, n = 4 mice. Bar values represent the mean ± SEM. ***p < 0.001; two-way ANOVA corrected with Bonferroni post-hoc test. (**d**) Photomicrograph of the expression of GCaMP7b (green) in the SCN in both control (scr, left) and experimental (shCdk5, right) animals. The red hatched oval indicates the placement of the optic fiber. Blue: Dapi, green: GFP (produced by jGCaMPP7). Scale bar 50 µm.

We observed an increase in calcium activity in control mice (scramble) during the second half of the 15 minutes of the light pulse at ZT14, which was also sustained for over 15 minutes after the light pulse (Fig. 4b, black trace). In contrast, the ΔF/F_0_ in shCdk5 mice during the second half and after the 15-minute light pulse was significantly lower compared to the control animals (Fig. 4b, red trace). This calcium activity was significantly decreased in shCdk5 mice during the last five minutes of the light pulse as compared to the baseline levels (see Methods; Fig. 4c; Supplementary Fig. 4d). Finally, mice were sacrificed, and the GFP signal was assessed by immunostaining to verify virus expression in the SCN (Fig. 4d). The outlined circle in red indicates where the fibers were located. Taken together, these results indicate that Cdk5 modulates Ca^2+^ mediated neuronal signaling.

### Cdk5 regulates the DARPP32-PKA axis

The cAMP-activated Protein Kinase A (PKA) signaling pathway, which leads to phosphorylation of CREB, plays a pivotal role in regulating phase delays in photic resetting ^50, 51^. Since the PKA signaling pathway can be induced *in vivo* ^10, 52^ and *in vitro* ^53^, we investigated whether Cdk5 could play a role in PKA-mediated CREB phosphorylation. To this end, we employed Förster resonance energy transfer (FRET), a widely used method to investigate molecular interactions between proteins such as CREB and CBP in living cells ^15, 54^.

We transfected control (wt) and *Cdk5* knock-out (*Cdk5* KO) NIH 3T3 cell lines ^34^ with ICAP (an indicator of CREB activation due to phosphorylation) and stimulated the cells with forskolin in the presence of Ca^2+^. The difference in phosphorylation before and after forskolin treatment of the CREB domain in the reporter decreases the FRET signal normally between 10 and 30 minutes, while no difference in phosphorylation brings the FRET signal back towards baseline. We observed that the FRET signal in control cells strongly decreased between 10 and 30 minutes after the stimulus compared to baseline (Fig. 5a, blue trace, the first 5 minutes are ignored, because they represent the diffraction of the solvent DMSO). In contrast, the FRET signal in *Cdk5* KO cells rose towards baseline after an initial decline in response to forskolin (Fig. 5a, red trace). This indicated that Cdk5 is involved in the phosphorylation of CREB. Notably, the forskolin solvent DMSO can’t stimulate CREB phosphorylation on its own (Supplementary Fig. 5a).

**Figure 5.**
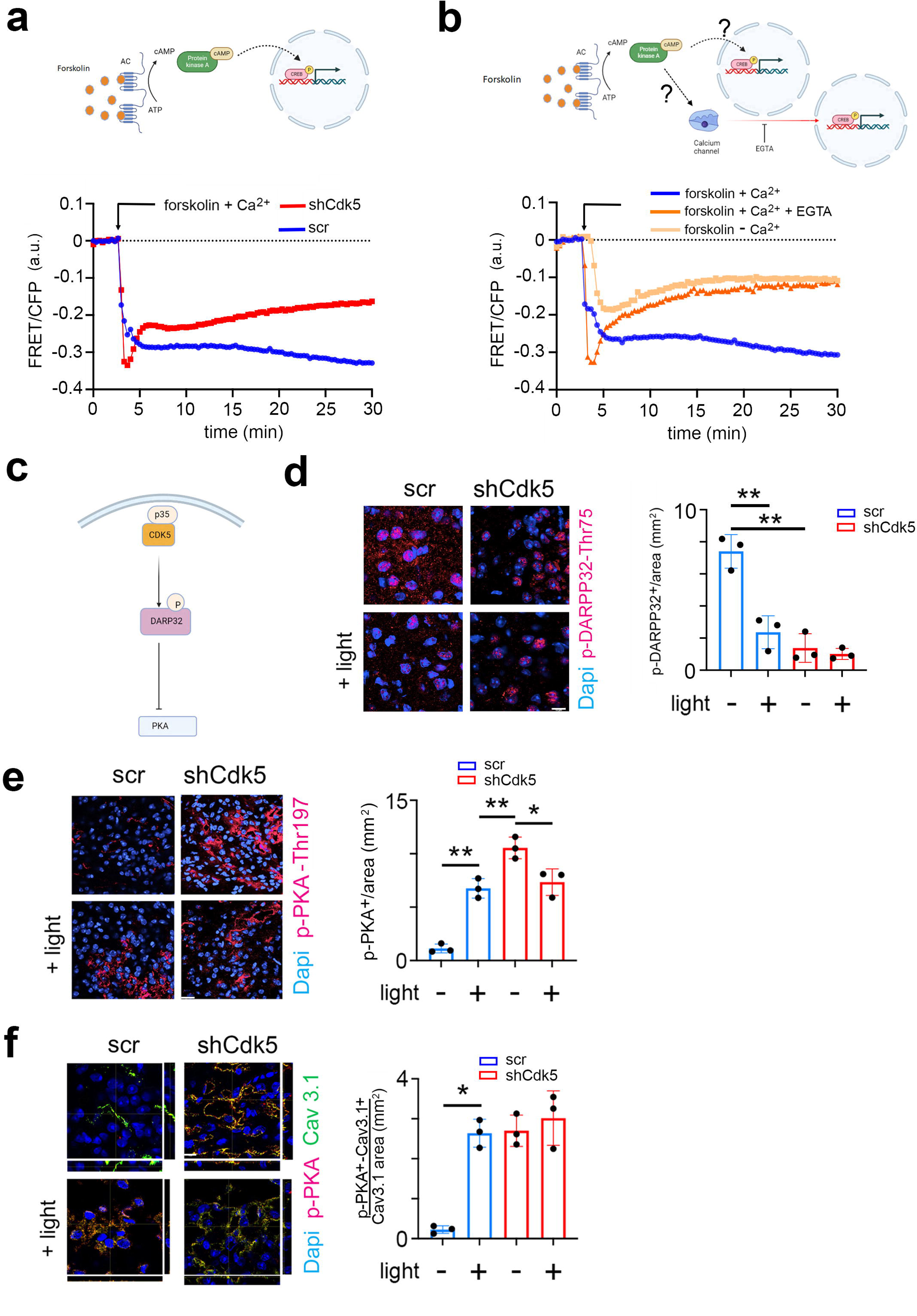
CDK5 regulates PKA phosphorylation via DARPP32 phosphorylation. (**a**) Top: Scheme of the forskolin-PKA-CREB signaling pathway. Bottom: FRET/CFP signal ratio changes in response to forskolin treatment in NIH 3T3 cells transfected with either a scr control (blue) or shCdk5 (red) expression construct. Values are the mean ± SD. Two-way ANOVA revealed a significant difference between the curves, n = 3, ****p < 0.0001. (**b**) Top: Scheme of the forskolin-PKA-CREB signaling pathway and calcium signaling. Bottom: FRET/CFP signal ratio changes in response to forskolin treatment in NIH 3T3 cells with the addition of Ca^2+^ (blue), without the addition of Ca^2+^ (salmon colored), and with the addition of Ca^2+^ and EGTA (orange). Values are the mean ± SD. Two-way ANOVA revealed a significant difference between the grey and blue/orange curves, n = 3, ****p < 0.0001. (**c**) Scheme of CDK5-DARPP32-PKA pathway. (**d**) Immunohistochemistry on the SCN of control (scr) and shCdk5 mice using an antibody recognizing phosphorylated Thr-75 of DARPP32 (p-DARPP32) before and after a light pulse at ZT14. The red color shows p-DARPP32 and the blue color represents Dapi stained nuclei of SCN cells. Scale bar: 10 µm. The right panel shows the quantification of the p-DARPP32 signal. Values are the mean ± SEM. Unpaired t-test with Welch’s correction, n = 3, **p < 0.01. (**e**) Immunohistochemistry on the SCN of control (scr) and shCdk5 mice using an antibody recognizing phosphorylated Thr-197 of PKA (p-PKA) before and after a light pulse at ZT14. The red color shows p-PKA and the blue color represents Dapi stained nuclei of SCN cells. Scale bar: 20 µm. The right panel shows the quantification of the p-PKA signal. Values are the mean ± SEM. Unpaired t-test with Welch’s correction, n = 3, *p < 0.05, **p < 0.01. (**f**) Immunohistochemistry on the SCN of control (scr) and shCdk5 mice using an antibody recognizing phosphorylated Thr-197 of PKA (p-PKA) and Cav3.1 before and after a light pulse at ZT14. The red color shows p-PKA, the green color Cav3.1, and the blue color represents Dapi-stained nuclei of SCN cells. The yellow color signifies the co-localization of PKA and Cav3.1. The stripes on the left and bottom of each micrograph show the z-stacks to confirm co-localization. Scale bar: 10 µm. The right panel shows the quantification of relative p-PKA/Cav3.1. Values are the mean ± SEM. Unpaired t-test with Welch’s correction, n = 3, *p < 0.05.

Previous studies have described that Ca^2+^-mediated CREB transcription of target genes requires PKA activity ^42^. However, it is not clear whether there is a parallel (synergistic) relationship between PKA and Ca^2+^ signaling pathways or whether they are sequentially dependent on each other (Fig. 5b, cartoon model). To address this question, we performed the following FRET experiment. NIH 3T3 cells were stimulated with forskolin in the presence of Ca^2+^ with EGTA (Ca^2+^ chelator) (Fig. 5b, red line), without EGTA (Fig. 5b, blue line) or completely depleted of Ca^2+^ (Fig. 5b, orange line). We observed that under normal conditions the FRET signal decreased, comparable to the signal seen in figure 5a, indicating higher Ser-133 KID phosphorylation compared to the baseline (Fig. 5b, blue line). When we added EGTA (removing Ca^2+^), the FRET signal increased to the baseline level after forskolin treatment (Fig. 5b, red line). The cells depleted of Ca^2+^ were also not responsive to the forskolin stimulus, as the FRET signal moved towards the baseline level within 30 minutes (Fig. 5b, orange line). Together, our results indicate that CREB phosphorylation is modulated by Cdk5 via Ca^2+^ signaling, as suggested in figure 3. Interestingly, PKA did not appear to directly phosphorylate CREB, as CREB did not pull-down p-PKA in an immunoprecipitation experiment. In contrast, p-CaMKIV did interact with CREB (Supplementary Fig. 5b, c), suggesting that CREB is most likely phosphorylated by CaMKIV, which is probably indirectly regulated by PKA activity.

Next, we aimed to investigate what the possible pathway could be through which PKA regulates CaMKIV. Previous studies have shown that Cdk5 regulates PKA activity via DARPP32 ^55^ (Fig. 5c). Therefore, we asked whether DARPP32 phosphorylation was light-dependent and whether Cdk5 would modulate this process. We sacrificed mice either receiving a light pulse at ZT14 or no light. Cryo-sections containing the SCN were stained with an antibody recognizing phosphorylated Thr-75 (pThr-75) of DARPP32. We observed that DARPP32 is highly phosphorylated at ZT14, with the light signal significantly reducing the phosphorylation levels in the cytoplasm and nuclei (Fig. 5d, blue bars; Supplementary Fig. 5d, e). In contrast, silencing of *Cdk5* led to a dramatic decrease in the pThr-75 signal in the cytoplasm and nuclei of SCN cells at ZT14, and light did not have an effect (Fig. 5d, red bars, Supplementary Fig. 5e). These observations are consistent with the view that Cdk5 phosphorylates DARPP32 and that light inhibits this process.

Non phosphorylated DARPP32 promotes PKA activity, characterized by phosphorylation at Thr-197 in the catalytic site of PKA ^56, 57^. Therefore, we asked whether decreased levels of p-DARPP32 after the light stimulus at ZT14 could inversely correlate with the phosphorylation state of PKA. We performed immunostaining on coronal brain sections containing the SCN using an antibody recognizing the phosphorylated Thr-197 of PKA. We observed that PKA phosphorylation significantly increased after the light pulse in the SCN tissue obtained from control (scr) mice (Fig. 5e, right panel, blue bars). However, in SCN from shCdk5 mice, the phosphorylation level was already elevated before the light pulse compared to scr control (Fig. 5e, left panels, top micrographs), and it was also sustained after the light pulse (Fig. 5e, left panels, bottom micrographs, right panel, red bars). Our results indicate that Cdk5 gates PKA phosphorylation induced by the light pulse at ZT14. Many observations indicate that active PKA can stimulate the Ca^2+^ influx through Cav3 T-type voltage-gated channels, including Cav3.1 ^27, 58^. The molecular mechanism normally requires physical interaction between the channel and PKA, followed by phosphorylation, which influences the gating properties ^59^. Therefore, we performed a co-immunostaining in the same SCN sections collected before (Fig. 3) to detect both Cav3.1 and phospho-PKA (the active form). We observed that the colocalization between Cav3.1 and phospho-PKA dramatically increased after the light pulse in the SCN tissue of control (scr) mice (Fig. 5f, scr left panels yellow color, and blue bars in the right panel). Interestingly, the colocalization level of the two proteins was already high in the shCdk5 SCN tissue before the light pulse, compared to controls (Fig. 5f scramble vs. shCdk5, left panel, top micrographs). The colocalization level between Cav3.1 and phospho-PKA in the shCdk5 tissues was not influenced by the light pulse (Fig. 5f, right panel, red bars). Altogether our results suggest that Cdk5 gates the PKA-Cav3.1 interaction in response to the light signal at ZT14 in an indirect way via DARPP32.

### Cdk5 affects light-induced gene expression

Light perceived in the dark period leads not only to phase shifts but also induces immediate early genes and certain clock genes in the SCN ^16, 17, 18, 60, 61^. This process involves the PKA – CaMK – CREB signaling pathway (reviewed in ^62^). Therefore, we investigated whether Cdk5 is involved in the signal transduction process to induce immediate early genes and clock genes in the SCN in response to light. To this end, we performed a time-course profile of light-induced genes and immediate early genes. We collected SCN from mice that received a nocturnal light pulse at ZT14 at different time points over 2 hours (Fig. 6).

**Figure 6.**
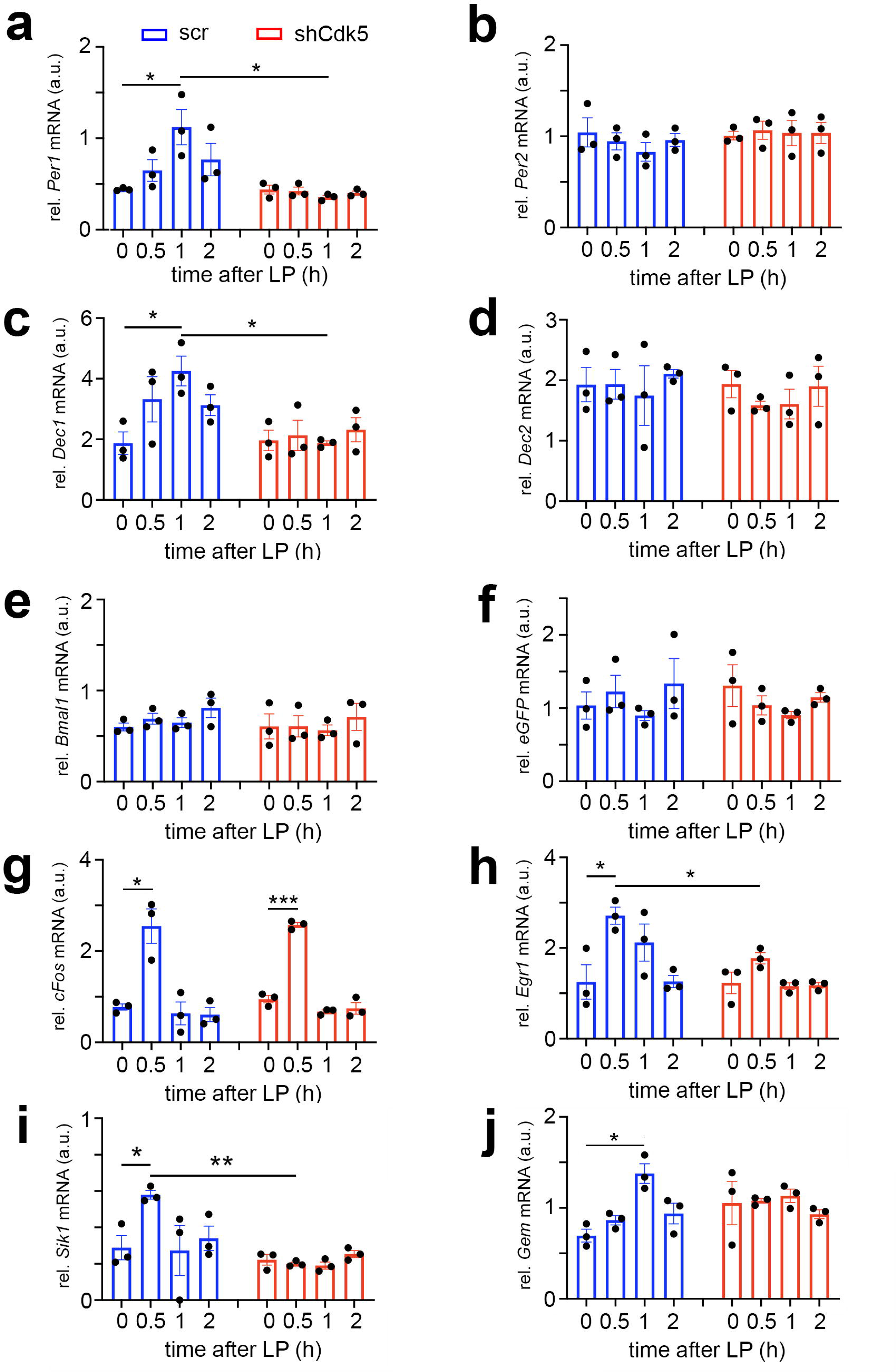
Cdk5 regulates light-induced gene expression in the SCN of some clock genes. Relative mRNA values are represented as blue bars for scr control animals and as red bars for shCdk5 mice. The values were determined 0, 0.5, 1, and 2 hours after a light pulse (LP) given at ZT14. (**a**) Induction of *Per1* mRNA expression by light with a maximum of 1 hour after light in scr control animals. In contrast *Per1* is not induced in shCdk5 SCN. Scr: 0 h: 0.44 ± 0.01, 0.5 h: 0.65 ± 0.12, 1 h: 1.12 ± 0.19, 2 h: 0.44 ± 0.01; shCdk5: 0 h: 0.44 ± 0.01, 0.5 h: 0.44 ± 0.01, 1 h: 0.44 ± 0.01, 2 h: 0.77 ± 0.18. Values are the mean ± SEM. Unpaired t-test, n = 3, *p < 0.05. (**b**) *Per2* mRNA expression is not induced by light neither in scr controls nor in shCdk5 animals. Scr: 0 h: 1.04 ± 0.16, 0.5 h: 0.95 ± 0.10, 1 h: 0.83 ± 0.10, 2 h: 0.96 ± 0.07; shCdk5: 0 h: 1.01 ± 0.05, 0.5 h: 1.07 ± 0.10, 1 h: 1.04 ± 0.14, 2 h: 1.04 ± 0.12. Values are the mean ± SEM. Unpaired t-test, n = 3 (**c**) Induction of *Dec1* mRNA expression by light with a maximum at 1 hour after light in scr control animals. In contrast *Dec1* is not induced in shCdk5 SCN. Scr: 0 h: 1.87 ± 0.37, 0.5 h: 3.32 ± 0.75, 1 h: 4.25 ± 0.49, 2 h: 3.13 ± 0.34; shCdk5: 0 h: 1.96 ± 0.34, 0.5 h: 2.13 ± 0.51, 1 h: 1.87 ± 0.07, 2 h: 2.32 ± 0.40. Values are the mean ± SEM. Unpaired t-test, n = 3, *p < 0.05. (**d**) *Dec2* mRNA expression is not induced by light neither in scr controls nor in shCdk5 animals. Scr: 0 h: 1.93 ± 0.29, 0.5 h: 1.94 ± 0.25, 1 h: 1.75 ± 0.49, 2 h: 2.11 ± 0.07; shCdk5: 0 h: 1.94 ± 0.23, 0.5 h: 1.58 ± 0.07, 1 h: 1.61 ± 0.25, 2 h: 1.90 ± 0.33. Values are the mean ± SEM. Unpaired t-test, n = 3. (**e**) *Bmal1* mRNA expression is not induced by light in the SCN of scr control and shCdk5 animals. Scr: 0 h: 0.60 ± 0.04, 0.5 h: 0.69 ± 0.06, 1 h: 0.65 ± 0.05, 2 h: 0.81 ± 0.12; shCdk5: 0 h: 0.61 ± 0.14, 0.5 h: 0.61 ± 0.12, 1 h: 0.56 ± 0.06, 2 h: 0.71 ± 0.15. Values are the mean ± SEM. Unpaired t-test, n = 3. (**f**) *eGFP* mRNA expression is detected in the SCN of scr control and shCdk5 animals demonstrating proper injection of expression constructs (scr: ssAAV-9/2-hSyn1-chl[1x(shNS)]-EGFP-WPRE-SV40p(A), shCdk5: ssAAV-9/2-hSyn1-chl[mouse(shCdk5)]-EGFP-WPRE-SV40p(A)). Scr: 0 h: 1.04 ± 0.19, 0.5 h: 1.23 ± 0.22, 1 h: 0.90 ± 0.07, 2 h: 1.34 ± 0.34; shCdk5: 0 h: 1.31 ± 0.28, 0.5 h: 1.04 ± 0.13, 1 h: 0.90 ± 0.05, 2 h: 1.15 ± 0.06. Values are the mean ± SEM. Unpaired t-test, n = 3. (**g**) Induction of *cFos* mRNA 0.5 hours after the light pulse in both scr controls and shCdk5 SCN. Scr: 0 h: 0.77 ± 0.07, 0.5 h: 2.55 ± 0.38, 1 h: 0.64 ± 0.25, 2 h: 0.61 ± 0.15; shCdk5: 0 h: 0.95 ± 0.08, 0.5 h: 2.57 ± 0.05, 1 h: 0.68 ± 0.04, 2 h: 0.74 ± 0.12. Values are the mean ± SEM. Unpaired t-test, n = 3, *p < 0.05, ***p < 0.001. (**h**) Induction of *Egr1* mRNA 0.5 hours after the light pulse in scr control but not shCdk5 SCN. Scr: 0 h: 1.25 ± 0.38, 0.5 h: 2.71 ± 0.19, 1 h: 2.12 ± 0.41, 2 h: 1.26 ± 0.13; shCdk5: 0 h: 1.23 ± 0.24, 0.5 h: 1.77 ± 0.13, 1 h: 1.16 ± 0.08, 2 h: 1.18 ± 0.06. Values are the mean ± SEM. Unpaired t-test, n = 3, *p < 0.05. (**i**) *Sik1* mRNA expression is induced by light in the SCN of scr control but not shCdk5 animals. Scr: 0 h: 0.29 ± 0.07, 0.5 h: 0.58 ± 0.02, 1 h: 0.27 ± 0.14, 2 h: 0.34 ± 0.07; shCdk5: 0 h: 0.22 ± 0.03, 0.5 h: 0.20 ± 0.01, 1 h: 0.19 ± 0.02, 2 h: 0.25 ± 0.02. Values are the mean ± SEM. Unpaired t-test, n = 3, *p < 0.05, **p < 0.01. (**j**) *Gem* mRNA expression is induced by light in the SCN of scr control but not shCdk5 animals. Scr: 0 h: 0.70 ± 0.07, 0.5 h: 0.86 ± 0.05, 1 h: 1.38 ± 0.11, 2 h: 0.94 ± 0.11; shCdk5: 0 h: 1.05 ± 0.24, 0.5 h: 1.08 ± 0.03, 1 h: 1.13 ± 0.07, 2 h: 0.93 ± 0.05. Values are the mean ± SEM. Unpaired t-test, n = 3, *p < 0.05.

In agreement with previous studies, *Per1* and *Dec1* mRNA expression was induced by light, peaking at 1hour after the stimulus. Conversely, *Per2* and *Dec2* mRNA expression was not affected by the light pulse at ZT14 (Fig. 6a-d, blue bars) ^18, 61, 63^. Knock-down of *Cdk5* abolished this light-driven *Per1* and *Dec1* gene induction (Fig. 6a, c, red bars), indicating the involvement of *Cdk5* in the light-driven activation process of these clock genes. As previously reported, expression of the clock gene *Bmal1* was not light-inducible ^34, 64^ and was not affected by shCdk5 (Fig. 6e). The injection of the control scr and shCdk5 constructs was successful, as demonstrated by the expression of *eGFP* mRNA in the analyzed SCN (Fig. 6f).

Interestingly, the knock-down of *Cdk5* did not affect light-mediated induction of *cFos* expression, which peaked at 0.5 hours after the light pulse (Fig. 6g). In contrast, *Egr1*, another immediate early gene involved in synaptic plasticity, learning, and memory ^65^, was light-inducible in control but not in shCdk5 animals (Fig. 6h). This suggests that the immediate early gene *cFos* is regulated by a different mechanism compared to *Egr1* and the clock genes *Per1* and *Dec1* in response to a light stimulus at ZT14.

Vasoactive intestinal polypeptide (VIP) has been described to play a role in phase-shifting the SCN clock ^66^. Furthermore, the light-induced expression of clock genes is localized in VIP-positive cells in the SCN, which are essential for clock resetting ^67^. Therefore, we tested whether *Vip* gene expression is affected by shCdk5. We observed that a light pulse did not significantly induce *Vip* expression in the SCN, nor did shCdk5 affect its general expression (Supplementary Fig. 6a). This suggests that *Cdk5* does not regulate *Vip* expression and modulate phase shifts via VIP.

Salt inducible kinase 1 (*Sik1*) is involved in the regulation of the magnitude and duration of phase shifts by acting as a suppressor of the effects of light on the clock ^68^. Therefore, we tested how a light pulse affected *Sik1* expression in the SCN and whether *Cdk5* might play a role in its regulation. We observed that *Sik1* was significantly induced by a light pulse in the SCN of control mice after 0.5 hours. However, the knock-down of *Cdk5* abolished this induction (Fig. 6i). This suggests that *Cdk5* modulates *Sik1* expression to regulate the magnitude of the behavioral response to light.

The light-inducible small G-protein Gem limits the circadian clock phase-shift magnitude by inhibiting voltage-dependent calcium channels ^69^. We tested whether a light pulse affected *Gem* expression in the SCN and whether this involved Cdk5. We observed that *Gem* was significantly induced by light 1 hour after light administration (Fig. 6j, blue bars). Interestingly, knock-down of *Cdk5* abolished this induction (Fig. 6i, red bars), but *Gem* levels seemed to be slightly elevated already before light administration (Fig. 6i, time point 0). This indicates that Cdk5 influences light induced *Gem* expression and may also affect basal *Gem* expression before the light pulse. Similar results for light induced gene expression in shCdk5 SCN were observed in SCN of *Per2^Brdm1^* mutant mice (Supplementary Fig. 6b, c).

Phase shifts of the circadian clock can also be studied in cell cultures using forskolin instead of light as a stimulus ^53^. In accordance with our *in vivo* experiments (Fig. 6), expression of *Per1* but not *Per2* mRNA was induced in synchronized NIH 3T3 fibroblast cells after forskolin treatment (Supplementary Fig. 6d, e, blue bars). Comparable to the experiments in the SCN, *Per1* induction was abolished in *Cdk5* knock-out cells (Supplementary Fig. 6d). In contrast*, cFos* mRNA induction was not affected in *Cdk5* knock-out cells (Supplementary Fig. 6f), consistent with our observations in the SCN (Fig. 6g).

Collectively, our expression data provide evidence that *Cdk5* regulates light- and forskolin-mediated expression of genes critical for the regulation of phase delays of the circadian clock. Immediate early genes, such as *Egr1*, are regulated in a similar manner, whereas others, such as *cFos*, are regulated by a different mechanism not involving *Cdk5*.

## Discussion

In this study, we investigated the role of Cdk5 in rapid phase shifts of the circadian clock. We found that Cdk5 activity is regulated by light and that Cdk5 is necessary for phase delays but not phase advances. We identified Cdk5 to play a major role in the modulation of Ca^2+^ levels and gating of the PKA-CaMK-CREB signaling pathway, coordinating it with the presence of PER2 in the nucleus of SCN cells.

In a previous study, we identified the protein kinase Cdk5 to regulate the phosphorylation and nuclear localization of the clock protein PER2 ^34^. Because PER2 and protein kinases are involved in the photic signaling mechanism of clock phase adaptation ^15, 20, 62, 70, 71^, we tested the involvement of Cdk5 in this process. The phenotype of *Cdk5* knock-down (shCdk5) in the SCN of mice resembled the phenotypes observed in *Per2* mutant (*Per2^Brdm1^*) and neuronal *Per2* knock-out (n*Per2* ko) mice. ShCdk5, as well as *Per2^Brdm1^* and n*Per2* ko animals, showed strongly reduced phase delays in response to a short light pulse given at ZT14 (Fig. 1a, c) ^20^ or CT14 (Fig. 1d, f) ^21, 72^. These mouse lines displayed a shortened period consistent with previous observations (Fig. 1b, e) ^34, 73^. Our results indicate that Cdk5 is not only involved in the regulation of the circadian clock mechanism via nuclear localization of PER2 but also plays an important role in the molecular mechanism that leads to a delay of clock phase in response to a light pulse in the early dark phase or early subjective night.

Since Cdk5 mediates the effects of light at the behavioral (Fig. 1) level, we tested the influence of light on Cdk5 protein accumulation and kinase activity in the SCN at ZT14 (Fig. 2). We observed no change in the protein accumulation of Cdk5. On the other hand, Cdk5 kinase activity was reduced in the SCN after a light pulse at ZT14 (Fig. 2d, e), which was surprising in the context of increased p35 levels (Fig. 2a, c) and augmented PKA phosphorylation (Fig. 2a, b). However, this observation is in line with what we previously reported, where we demonstrated that Cdk5 kinase activity was low during the light phase and higher during the dark phase ^34^. It appeared, however, that p35 was not interacting with Cdk5 after light at ZT14 (Fig. 2f). Additional interactions of Cdk5 with unknown proteins may also be lost (Supplementary Fig. 2e). These observations suggest that Cdk5 was most likely modified in response to light leading to loss of interaction with p35 and other proteins. Ser159 of Cdk5 mediates the specificity of the Cdk5-p35 interaction ^74^, and therefore, phosphorylation of this site by an unknown kinase may mediate the loss of Cdk5 activity. Several additional phosphorylation sites in Cdk5 have been identified, of which phosphorylation of S47 renders Cdk5 inactive ^75^. Which one of the phosphorylation sites in Cdk5 is modulated by light and what additional interactors may be involved in this process remains to be established.

Light in the early portion of the dark phase elicits phase delays, which involve T-type calcium channels, PKA-signaling, and Ca^2+^ signaling, ending in the phosphorylation of CREB (reviewed in ^62^). We observed that in shCdk5 mice, CREB was already phosphorylated in the absence of light, although the total protein amount did not change (Fig. 3a, Supplementary Fig. 3a, b). Similarly, CaMKII and CaMKIV were shown to be phosphorylated and, therefore, activated only after the light pulse in control animals (Fig. 3b, d). Conversely, these kinases were highly phosphorylated in a light-independent manner in the SCN of shCdk5 animals (Fig. 3b, d), indicating that Cdk5 had a suppressive function on the phosphorylation of CaMKII and CaMKIV.

A stimulus can promote calmodulin (CAM) involving CaMKII gamma to translocate from calcium channels to the nucleus to promote CaMKIV phosphorylation and activation ^44^. Unexpectedly, we observed that a light stimulus can have a similar but distinct effect on CAM in SCN cells (Fig. 3c). CaMKII alpha was phosphorylated after a light pulse at ZT14, which led to perinuclear localization of CAM in control mice, while this localization pattern was already observed in shCdk5 animals independently of the light stimulus (Fig. 3c). In control mice that received no light, CAM showed a diffuse expression pattern similar to the T-type calcium channel Cav3.1 at ZT14.

Interestingly, this light-driven localization pattern was echoed by the change in cellular distribution of the T-type calcium channel Cav3.1 known as internalization/externalization. Again, the presence of Cdk5 suppressed the localization of this channel to the cell membrane in the absence of light, with light allowing localization to the cell membrane (Fig. 3e). This observation is reminiscent of investigations described previously in which Cdk5 appeared to play an important role in channel translocation ^76, 77^ as well as in receptor translocation ^46, 47^. Thus, our findings are in accordance with the view that Cdk5 plays a crucial role in light stimulus driven cell dynamics.

Calcium plays an important role in circadian and phase-shifting biology ^48^. Circadian calcium fluxes in the cytosol of SCN neurons have been demonstrated *in vitro* ^78^, and they change rapidly as a response to light perceived by the retina ^49^. We performed *in vivo* live imaging to detect Ca^2+^ levels in the SCN using fiber photometry with protein-based Ca^2+^ indicators such as GCaMP ^79^. With this approach, we observed that calcium fluxes in the SCN of control mice increased during and after a light pulse, but this change was significantly dampened in shCdk5 animals (Fig. 4b, c). Interestingly, although the Ca^2+^ influx was generally reduced in the SCN of shCdk5 mice, we observed random Ca^2+^ activity, which was independent of any light stimulus. These transients were observed also at the beginning of ZT14, before the light pulse (Fig. 4b, c). These results may indicate the presence of a calcium leak reminiscent of the already active phosphorylation cascade observed in the shCdk5 SCN in the absence of light (Fig. 3). We do not know, however, whether internal calcium stores involving ryanodine receptors ^80^ are altered by Cdk5 as well and how this would contribute to the observed phenotypes.

The PKA signaling pathway is involved in the resetting of the circadian phase (reviewed in ^50^). Interference with PKA activation in the early subjective night led to reduced phase delay responses as observed *in vitro* in the SCN ^52^. Here we find that Cdk5 plays an inhibitory role in PKA phosphorylation and activation. The FRET approach shows that in cells the lack of Cdk5 makes cells unresponsive to forskolin (Fig. 5a), an agent known to mitigate phase shifts in cells via PKA ^53^. Interestingly, PKA appears to influence phase shifts and CREB phosphorylation indirectly via a Ca^2+^ dependent mechanism (Fig. 5b) with phosphorylated CaMKIV being the kinase that phosphorylates CREB (Supplementary Fig. 5b, c). This observation is in contrast with previous studies that suggested a direct phosphorylation of CREB by PKA ^10, 52^. However, p-PKA is mostly located in the cytoplasm (Fig. 5e) while p-CaMKIV is in the nuclei (Fig. 3d). Furthermore, our experiments indicate that CREB did not interact with p-PKA but did with p-CaMKIV (suppl Fig. 5b, c), supporting the notion that PKA regulates CREB phosphorylation indirectly via CaMKIV in the SCN.

Because we observed that PKA was already phosphorylated in the dark when Cdk5 was silenced (Fig.5e), we asked how Cdk5 could negatively regulate PKA phosphorylation. A previous study described that Cdk5 can phosphorylate DARPP32 to suppress PKA activity ^81^. Furthermore, *Darpp-32* KO mice show attenuated phase delays ^12^ resembling shCdk5 mice (Fig. 1). In accordance with these studies, we found an inverse correlation between p-DARPP32 (Fig. 5d) and p-PKA (Fig. 5e), implying that Cdk5 indirectly inhibits PKA activity via DARPP32. However, phosphatases such as PP2A and calcineurin, which de-phosphorylate DARPP32 including the Cdk5 phosphorylation site, may be involved in this process as well ^82^. Upon light treatment and increase of Ca^2+^, these phosphatases would dephosphorylate DARPP32 and thereby inactivate it, leading to PKA activation. This process may occur in parallel to the Cdk5 regulation of DARPP32 contributing to a sustained activation of the light signaling pathway via PKA activation.

Our results imply that PKA action on CREB might be mediated via T-type calcium channels such as Cav 3.1 (Fig. 5f). This assumption is reasonable because PKA can phosphorylate Cav 3.1 channels and increase electrical conductivity, which leads to a higher influx of Ca^2+ 26^. To that extent, our results indicate that a higher co-localization of p-PKA with Cav 3.1 is associated with an activation of the CaMK pathway and CREB phosphorylation.

Light-induced phosphorylation of CREB leads to induction of immediate early genes and clock genes (reviewed in ^62^). Accordingly, we observed that the clock genes *Per1* and *Dec1* but not *Per2*, *Dec2,* and *Bmal1* were induced in the SCN by light at ZT14 (Fig. 6a-e, blue bars) consistent with previous findings ^15, 18, 61, 63^. The light induction of *Per1* and *Dec1* was abolished in shCdk5 animals (Fig. 6a, c, red bars) as well as in *Per2^Brdm1^* mutant mice (Supplementary Fig. 6b, c), suggesting involvement of Cdk5 and Per2 in induction of these genes. In contrast, light induction of the immediate early gene *cFos* was neither affected in shCdk5 nor *Per2^Brdm1^* SCN (Fig. 6g, Supplementary Fig. 6f), resembling the normal *cFos* induction in *Per2* KO animals ^15^. This indicates that the light signaling mechanism for *cFos* induction is different from the one mediating induction of *Per1* and *Dec1*. Interestingly, however, the light-inducible genes *Sik1* and *Gem,* which are involved in limiting the effects of light on the clock ^68, 69^ were not light-inducible in shCdk5 animals (Fig. 6i, j) supporting the view that the factors that drive (*Per1, Dec1*) or limit (*Sik1, Gem*) the effects of light on the clock are regulated by the same mechanism. Interestingly, neither lack of *Per1*, *Dec1* ^20, 83^ nor *Sik1* or *Gem* ^68, 69^ alone abolish phase delays. Of note is that lack of *cFos* or *Egr1* did not affect phase delays either ^84, 85^. Furthermore, the neuropeptide vasoactive intestinal peptide (VIP), which is important in circadian light responses ^67^, was not inducible by a light pulse at ZT14 in control as well as shCdk5 animals (Supplementary Fig. 6a), indicating that Cdk5 acts upstream of VIP signaling. Overall, the present data suggest that Cdk5 not only regulates the light-sensitive PKA - CaMK-CREB signaling pathway but ultimately also affects gene expression. For the transcriptional activation of those genes, nuclear PER2 protein is necessary, which is regulated by Cdk5 ^15, 34^. The combination of lack of induction of many genes in the Cdk5-regulated pathway is responsible for the manifestation of rapid behavioral phase delays.

Based on this and our previous studies, we propose the following molecular model for light-mediated phase delays (Fig. 7). The model is divided into two parts. One part describes the state before the light pulse, and the second part the mechanism after the light pulse. The state before the light pulse (ZT12-14) is depicted in the gray area in Figure 7. As reported previously, Cdk5 is active right after dark onset ^34^, depicted as the active Cdk5/p35 complex (blue). This has two consequences: 1) PER2 (red) is phosphorylated and translocates to the nucleus ^34^, and 2) DARPP32 is phosphorylated and thus inhibiting PKA activity ^81^. Hence, before the light pulse at ZT14, the nucleus is supplied with PER2, which appears to be necessary for light-mediated behavioral phase delays ^20, 21, 72^. In parallel, the signaling pathway necessary to phosphorylate CREB is turned off. This state can then be dramatically changed when light is applied at ZT14 evoking glutamate and PACAP release at the synapses between the RHT and the SCN. Interaction between p35 and Cdk5 is abolished (Fig. 2) thereby inactivating Cdk5 and stopping phosphorylation of PER2 and DARPP32. Since significant amounts of PER2 are already in the nucleus this probably has no consequences on nuclear PER2 function. However, DARPP32 is not phosphorylated anymore and the block on PKA is released. At the same time, PKA becomes phosphorylated due to PACAP and cAMP signaling, leading to activation of Cav3.1 by PKA (Fig. 5f) ^26^. This results in CaMKII and CaMKIV phosphorylation and, ultimately, to the phosphorylation of CREB in the nucleus (Fig. 3) ^45^. Phospho-CREB builds up a complex with CRTC1/CBP and PER2 ^15^ to activate gene expression of the *Per1, Dec1, Sik1,* and *Gem* genes. In the activation complex the amount of PER2 present in the nucleus may at least in part affect the magnitude of the phase delay, which is depending on the time the light pulse is given. In conclusion, Cdk5 activity is gating both processes, the pre-light condition as well as the post-light condition, leading to a concerted activation of a set of light-responsive genes that impinge on behavioral phase delays in response to nocturnal light exposure.

**Figure 7.**
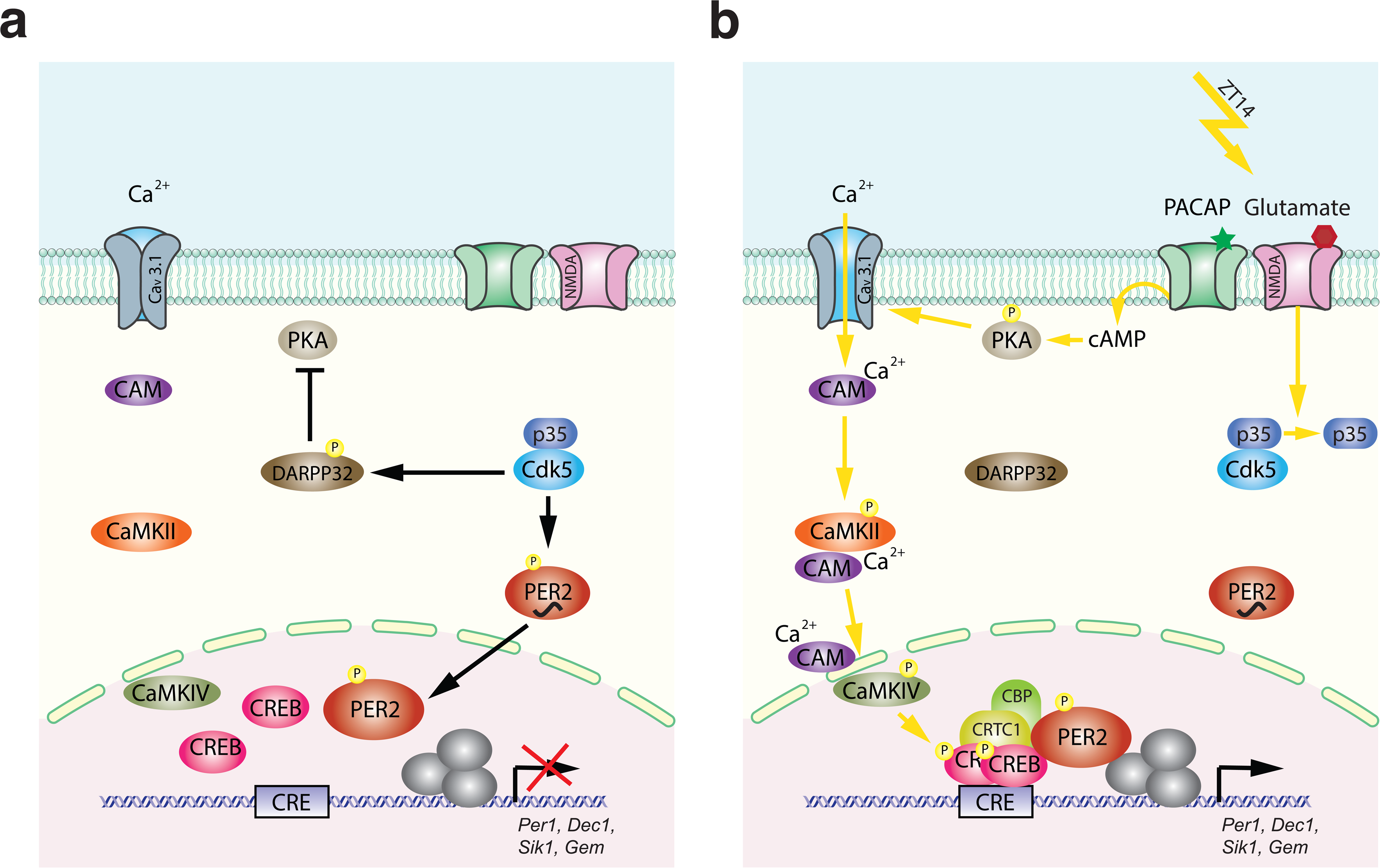
Model of Cdk5 gated light signal. (**a**) Cdk5 is active during the dark portion of the day. Active Cdk5 with its co-activator p35 phosphorylates PER2, which leads to stabilization and nuclear translocation of this protein that is abundant at ZT12. At the same time CDK5 phosphorylates DARPP32, which inhibits the PKA signaling pathway. (**b**) Light perceived in the dark phase at ZT14 leads to detachment of p35 from Cdk5 stopping Cdk5 activity. DARPP32 is not phosphorylated and hence can’t inhibit PKA. PKA that is activated by the light signal is phosphorylated and can mediate CREB phosphorylation via T-type calcium channels (Cav3.1) and the CaMK pathway leading to a transcriptionally active complex on the CRE element present in the promoters of many light responsive genes such as *Per1*, *Dec1*, *Gem* and *Sik1*. Overall, a light pulse at ZT14 will activate CREB phosphorylation and a protein complex will form. This complex needs phosphorylated PER2 that has accumulated in the nucleus between ZT12 and ZT14 to initiate the transcription of light-responsive genes. Both arms are necessary to build up a transcriptionally functional complex. Both arms depend on the presence and activity of CDK5, which therefore gates the light signal at ZT14. It is very likely that the amount of PER2 protein in the nucleus determines at least in part the magnitude of the phase delay, which depends on the timing of the light signal.

## Methods

### Animals and housing

All mice were housed with chow food (3432PX, Kliba-Nafag) and water ad libitum in transparent plastic cages (267 mm long, ×207 mm wide, ×140 mm high; Techniplast Makrolon type 2 1264C001) with a stainless-steel wire lid (Techniplast 1264C116), kept in soundproof ventilated chambers at constant temperature (22 ± 2 °C) and humidity (40 – 50%). All mice were entrained to a 12-h light–dark cycle (LD cycle), and the time of day was expressed as Zeitgeber time (ZT; ZT0 lights on, ZT12 lights off). Four-month-old 129/C57BL6 mixed males were used for the experiments. Housing and experimental procedures were performed per the guidelines of the Schweizer Tierschutzgesetz and the declaration of Helsinki. The state veterinarians of the Canton of Fribourg and Bern approved the protocol (license numbers: 2021-19-FR; BE45/18; BE21/22).

### Locomotor activity monitoring

Locomotor activity parameters were analyzed by monitoring wheel-running activity, as described in ^85^, and calculated using the ClockLab software (Actimetrics). To analyze free-running rhythms, animals were entrained to LD 12:12 and released into constant darkness (DD). The internal period length was determined from a regression line drawn through the activity onsets of ten days of stable rhythmicity under constant conditions, calculated using the respective inbuilt functions of the ClockLab software (Acquisition Version 3.208, Analysis Version 6.0.36). For better visualization of daily rhythms, locomotor activity records were double-plotted, which means that each day’s activity is plotted twice, to the right and below that of the previous day. For the analysis of light-induced resetting, we used Aschoff type II and I protocols ^86^. For type II, mice maintained in LD 12:12 were subjected to a 15 min. light pulse (LP, 500 lux) at ZT10 (no phase shift), 14 (phase delay), and 22 (phase advance). Subsequently, they were released into DD for ten days, and phase shift was measured. For type I, mice maintained in DD were subjected to a 15 min. light pulse at Circadian Time (CT) 10, 14, or 22. A circadian hour equals 1/24 of the endogenous period (τ), calculated as follows: circadian hour = tau/24 hours. To convert ZT hours to CT hours, we performed the following calculations:

- CT12 Day B - CT12 Day A + τ - 24 hrs
- CTX0-12 = CT12Day B – X* 1 circadian hour [X= CT12-CTx]
- CTX12-24 = CT12Day B + X * 1 circadian hour [X= CTx- CT12]

The phase shift was determined by fitting a regression line through the activity onsets of at least 7 days under LD conditions before the light pulse and a second line through the activity onsets of at least 7 days under DD after the light pulse. The first two days after the administration of the light pulse were not considered for the calculation of the phase shift. The distance between the two regression lines determined the phase shift. Before starting any new protocol, mice were allowed to stabilize their circadian oscillator for 10 days. The corresponding figure legends indicate the number of animals used in the behavioral studies.

### Light pulse and tissue isolation

Light pulse (LP., 500 lux) was given at ZT14, and mice were sacrificed at appropriate indicated times. Brains were collected, and SCN tissue was isolated for western blot or RT-qPCR use. For immunofluorescence experiments, mice were perfused with 4% PFA and cryoprotected in 30% sucrose. Tissue isolation at ZT14 without a light pulse was used as light-induction negative control.

### RNA extraction and cDNA synthesis

Total RNA was extracted from confluent 6 cm petri dishes or frozen SCN tissue using the Microspin RNA II kit (Machery & Nagel, Düren, Germany) according to the manufacturer’s instructions. 0.5 μg of total RNA was converted to single-strand cDNA in a total volume of 10 μL using the SuperScript IV VILO kit (Thermo Fisher Scientific, Waltham MA, USA) according to the manufacturer’s instructions. The samples were diluted to 200 μL with pure water. 5 μL of each sample was mixed with 7.5 μL of KAPA probe fast universal real-time PCR master mix (Merck, Darmstadt, Germany) and 2.5 μL of the indicated primer/probe combinations. For the subsequent real-time PCR, a Rotorgene 6000 machine was used (Qiagen, Hilden, Germany) and analyzed with the propriety software.

### qPCR primers

Per1:

FW: GGC ATG ATG CTG CTG ACC ACG RV: ACT GGG GCC ACC TCC AGT TC

TM: FAM-TGG CCC TCC CTC ACC TTA GCC TGT TCC T-BHQ1

Per2:

FW: TCC ACA GCT ACA CCA CCC CTT A RV: TTT CTC CTC CAT GCA CTC CTG A

TM: FAM-CCG CTG CAC ACA CTC CAG GGC G-BHQ1

Dec1

FW: TGC AGA CAG GAG CGC ACA GT RV: GCT TTGGGC AGG CAG GTA GGA

TM: FAM-TGG TTG CGC GCT GGG GAT CCG T-BHQ1

Dec2

FW: ACA GAA TGG GGA GCG CTC TCT GAA RV: TGA AAC CCC GAG TGG AAC GCA

TM: FAM-CGC CGG TCC AGG CCG ACT TGG A-BHQ1

Bmal1

FW: GCA ATG CAA TGT CCA GGA AG RV: GCT TCT GTG TAT GGG TTG GT

TM: FAM- ACC GTG CTA AGG ATG GCT GTT CAG CA-BHQ1 eGFP

FW: CAT CTG CAC CAC CGG CAA GC RV: GGT CGG GGT AGC GGC TGA A

TM: FAM- TGC CCG TGC CCT GGC CCA CC-BHQ1

cFos

FW: GCC GGG GAC AGC CTT TCC TA

RV: TCT GCG CAA AAG TCC TGT GTG TTG A

TM: FAM-CCA GCC GAC TCC TTC TCC AGC ATG GGC-BHQ1

Egr1

FW: CGG CAG CAG CGC CTT CAA T

RV: GGA CTC TGT GGT CAG GTG CTC AT

TM: FAM-CCT CAA GGG GAG CCG AGC GAA CAA CCC-BHQ1

Sik1

FW: GGC TGC ACG ACC AGC AAT CG

RV: GGC GGT AGA AGA GTG GTG CTG TA

TM: FAM- TCC TGC ACC AGC AGA GGC TGC TCC AG-BHQ1

Gem

FW: TGG GAA AAT AAG GGG GAG AA RV: AGC TTG CAC GGT CTG TGA TA

TM: FAM- CCA CTG CAT GCA GGT CGG GGA TGC C-BHQ1

Vip

FW: AGC AGA ACT TCA GCA CCC TAG ACA RV: TCG GTG CCT CCT TGG CTG TT

TM: FAM- AGC CGG AAA GGC AGC CCT GCC T-BHQ1

Tprkb (normalisation probe for Tprkb)

FW: GGC TGG CAT CAG ACC CAC AGA RV: GGG CCC GTA GAG TCG GGA AA

TM: FAM-CCT GCG TCT GCC CTC TGA GGG CTG-BHQ1

Atp5h (normalisation probe for Atp5h)

FW: TGC CCT GAA GAT TCC TGT GCC T RV: ACT CAG CAC AGC TCT TCA CAT CCT

TM: FAM-TCT CCT CCT GGT CCA CCA GGG CTG TGT-BHQ1

Sirt2 (normalisation probe for Sirt2)

FW: CAG GCC AGA CGG ACC CCT TC RV: AGG CCA CGT CCC TGT AAG CC

TM: FAM- TGA TGG GCC TGG GAG GTG GCA TGG A-BHQ1

Nono (normalisation probe for Nono)

FW: TCT TTT CTC GGG ACG GTG GAG RV: GTC TGC CTC GCA GTC CTC ACT

TM: FAM- CGT GCA GCG TCG CCC ATA CTC CGA GC-BHQ1

### Immunofluorescence

SCN cryosections (40 µM) were placed in a 24-well plate, washed three times with 1x TBS (0.1 M Tris/0.15 M NaCl) and 2x SSC (0.3 M NaCl/0.03 M tri-Na-citrate pH 7). Antigen retrieval was performed with 2xSSC heating to 85°C for 30 min. Then, sections were washed twice in 2x SSC and three times in 1x TBS pH 7.5 before blocking them for 1.5 hours in 10% fetal bovine serum (Gibco)/0.1% Triton X-100/1x TBS at RT. If the recipient species for some raised antibody was the mouse, we performed a Mouse on Mouse (MOM; Ab269452) blocking (2h) before 10% FBS to block endogenous mouse immunoglobulins in a mouse tissue section. After the blocking, the primary antibodies (Table 1), diluted in 1% FBS/0.1% Triton X-100/1x TBS, were added to the sections and incubated overnight at 4°C. The next day, sections were washed with 1x TBS and incubated with the appropriate fluorescent secondary antibodies diluted 1:500 in 1% FBS/0.1% Triton X-100/1x TBS for 3 hours at RT. (Alexa Fluor 488-AffiniPure Donkey Anti-Rabbit IgG (H+L) no. 711–545–152, Lot: 132876, Alexa Fluor647-AffiniPure Donkey Anti-Mouse IgG (H+L) no. 715–605–150, Lot: 131725, Alexa Fluor647-AffiniPure Donkey Anti-Rabbit IgG (H+L) no. 711–602–152, Lot: 136317 and all from Jackson Immuno Research). Tissue sections were stained with DAPI (1:5000 in PBS; Roche) for 15 min. Finally, the tissue sections were rewashed twice in 1x TBS and mounted on glass microscope slides. Fluorescent images were taken using a confocal microscope (Leica TCS SP5), and pictures were taken with a magnification of 63x with or without indicated additional zoom. Images were processed with the Leica Application Suite Advanced Fluorescence 2.7.3.9723. Immunostained sections were quantified using ImageJ version 1.49. Statistical analysis was performed on three animals per treatment.

**Table 1.** Antibodies used for the immunostainings.

| Antibody | Specie | Company | Catalog number | Dilution |
| --- | --- | --- | --- | --- |
| anti-Cdk5 clone 2H6 | Mouse | Origene | CF500397 | 1:100 |
| anti-GFP | Rabbit | Abcam | ab6556 | 1:500 |
| Anti-Creb | Rabbit | Cell signaling | D76D11 | 1:200 |
| Anti-Creb (pSer133) | Rabbit | Abcam | Ab32096 | 1:500 |
| Anti-CaMKII | Rabbit | Abcam | Ab52470 | 1:200 |
| Anti-CaMKII (pThr286) | Mouse | Invitrogen | MA1-047 | 1:100 |
| Anti-CaMKIV | Rabbit | Abcam | Ab3557 | 1:200 |
| Anti-CaMKIV (pThr196/200) | Rabbit | Invitrogen | PA5-105011 | 1:100 |
| Anti-CaV3.1 | Rabbit | Invitrogen | PA5-50635 | 1:100 |
| Anti-Calmodulin | Mouse | Invitrogen | MA3-917 | 1:100 |
| Anti-PKA (pT197) | Rabbit | Abcam | Ab75991 | 1:100 |
| Anti-Darpp32 (pThr75) | Rabbit | Invitrogen | PA5-105037 | 1:100 |

### Adeno Associated Virus (AAV) production and stereotaxic injections

Experiments were performed as previously described ^34^. Stereotaxic injections were performed on 4- to 5-month-old mice under isoflurane anesthesia using a stereotaxic apparatus (Stoelting). The brain was exposed by craniotomy, and the Bregma was used as a reference point for all coordinates. AAVs were injected bilaterally into the SCN (Bregma: anterior-posterior (AP) − 0.40 mm; medial-lateral (ML) ±0.00 mm; dorsal-ventral (DV) – 5.7 mm, angle + /- 3°) using a hydraulic manipulator (Narishige: MO-10 one-axis oil hydraulic micromanipulator, http://products.narishige-group.com/group1/MO-10/electro/english.html) at a rate of 40 nL/min through a pulled glass pipette (Drummond, 10 µl glass micropipette; Cat number: 5-000-1001-X10). The pipette was first raised 0.1 mm to allow the spread of the AAVs and later withdrawn 5 min after the end of the injection. After surgery, mice were allowed to recover for 2 weeks and entrained to LD 12:12 before behavior and molecular investigations.

The injected viruses were:

- SsAAV-9/2-hSyn1-chI[mouse(shCdk5)]-EGFP-WPRE-SV40p(A)
- ssAAV-9/2-hSyn1-chI[1x(shNS)]-EGFP-WPRE-SV40p(A)

### Protein extraction from SCN tissue

The protocol was a modified version of what was published before ^15^. Isolated SCNs obtained from 4 different mice were pooled according to the indicated condition (either dark or 15 min after the light pulse). The pooled tissues were frozen in liquid N2 and resuspended in a brain-specific lysis buffer (50 mM Tris-HCl pH 7.4, 150 mM NaCl, 0.25% SDS, 0.25% sodium deoxycholate, 1 mM EDTA). They were homogenized using a pellet pestle, kept on ice for 30 min and vortexed for 30 s, followed by N2 freezing. Frozen samples were left to melt on ice. The samples were sonicated (10 s, 15% amplitude) and centrifuged for 20 min at 16,000 g at 4 °C. The supernatant was collected in new tubes, and the pellet was discarded.

### Immunoprecipitation

The protocol was described before ^34^. The protein extract was diluted with the appropriate lysis buffer in a final volume of 250 µL and immunoprecipitated using the indicated antibody (ratio 1:50). The reaction was kept at 4°C overnight on a rotary shaker. The day after, samples were captured with 50 µL of 50% (w/v) protein-A agarose beads (Roche), and the reaction was kept at 4°C for 3 hr on a rotary shaker. Before use, beads were washed thrice with the appropriate protein buffer and resuspended in the same buffer (50% w/v). The beads were collected by centrifugation and washed three times with NP-40 buffer (100 mM Tris-HCl pH7.5, 150 mM NaCl, 2 mM EDTA, 0.1% NP-40). After the final wash, beads were resuspended in 2% SDS 10%, glycerol, 63 mM Trish-HCL pH 6.8, and proteins were eluted for 15 min at RT. Laemmli buffer was finally added, and samples were boiled for 5 min at 95° C and loaded onto 10% SDS-PAGE gels.

### Western blot

Circa 40 µg o of protein was loaded onto 10% SDS-PAGE gel and run at 100 Volt for two hours. Protein migration was followed by a semidry transfer (40 mA, 1 h 30 s) using Hybond® ECL™ nitrocellulose. We subsequently performed red ponceau staining (0.1% of Ponceau S dye and 5%) on the membrane to confirm the successful transfer. The list of antibodies used in the paper is shown in Table 2. The membrane was washed with TBS 1x/Tween 0.1% and blocked with TBS 1x/BSA 5%/Tween 0.1% for 1 h. After washing, the membrane was blotted with the appropriate primary antibodies overnight. The day after, membranes were washed three times with TBS 1x/Tween 0.1%, followed by secondary antibody immunoblotting for 1 h at room temperature. The densitometric signal was digitally acquired with the Azure Biosystem.

**Table 2.**
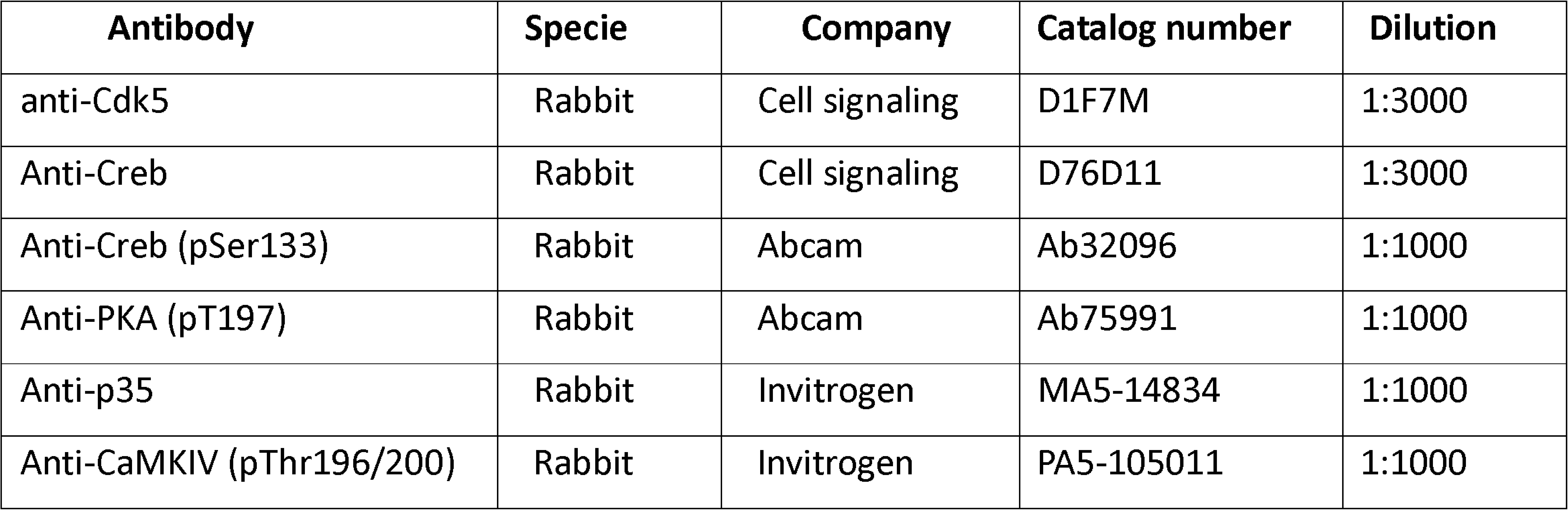
Antibodies used for the western blots.

All the original blots are shown in Supplementary Figure 7.

### *In vitro* kinase assay using immunoprecipitated Cdk5 from SCN

The protocol is the same as before ^34^. CDK5 was immunoprecipitated (4°C overnight with 2x Protein A agarose (Sigma-Aldrich)) from SCN samples at ZT14 in the dark or after a light pulse (LP., circa 500 lux) of 15min. After immunoprecipitation, samples were diluted in a washing buffer and split into two halves. One-half of the IP was used for an *in vitro* kinase assay. Briefly, 1 µg of histone H1 (Sigma-Aldrich) was added to the immunoprecipitated CDK5. Assays were carried out in reaction buffer (30 mM HEPES, pH 7.2, 10 mM MgCl_2_, and 1 mM DTT) containing [γ-^32^P] ATP (10 Ci) at 30°C for 1 hour and then terminated by adding SDS sample buffer and boiling for 5 min. Samples were subjected to 15% SDS-PAGE, stained by Coomassie Brilliant Blue, and dried, and then phosphorylated histone H1 was detected by autoradiography. The other half of the IP was used for Western blotting to determine the total CDK5 immunoprecipitated from the SCN samples. The following formula was used to quantify the kinase activity at each time point: ([^32^P] H1/total H1)/CDK5 ^IP protein^.

### Cell Culture

NIH3T3 wt and CRISPR/Cas9 *Cdk5* ko ^34^ mouse fibroblast cells (ATCCRCRL-1658) were maintained in Dulbecco’s modified Eagle’s medium (DMEM), containing 10% fetal calf serum (FCS) and 100 U/mL penicillin-streptomycin at 37°C in a humidified atmosphere containing 5% CO2. For any experiment, cells were synchronized with forskolin (100 µM).

### Surgical procedure for fiber photometry recordings

Animals previously infected either with AAV9-hSyn1-chI[1x(shNS)]-jGCaMP7b-WPRE-SV40p(A) (*scramble*) or AAV9-hSyn1-chI[mouse(shCdk5)]-jGCaMP7b -WPRE-SV40p(A) (*shCDK5*), were injected with Metacam (Meloxicam, 5mg/kg s.c.) for analgesia, then anesthetized with 1.5 –2 % isoflurane/O_2_ mix. Mice were placed in a Kopf digital stereotactic frame. Their body temperature was kept constant at 37 °C via a feedback-coupled heating device (Panlab/Harvard Apparatus), and their eyes were covered with ointment (Bepanthen Augen- und Nasensalbe, Bayer). After the skin incision (formerly prepared aseptically), the skull bone was cleaned with saline to remove the remaining tissue. A small craniotomy to target the SCN was drilled into the skull (Micro-Drill from Harvard Apparatus with burrs of 0.7 mm diameter from Fine Science Tools), and the dura was carefully removed. An optical fiber implant (400 μm, 0.5 NA Core Multimode Optical Fiber, FT400ERT, inserted into ceramic ferrules, 2.5 mm OD; 440 μm ID, Thorlabs) was slowly implanted above the SCN to allow for imaging of GCaMP7b signals (AP: +0.40; ML: ±0.0; DV: -5.3; angle +/- 4°). One stainless steel screw was inserted into the skull over the cerebellum for stability purposes. The implant was then secured to the skull with dental cement (Paladur, Kulzer). After surgical procedures, mice were allowed to recover for one week and finally tethered with an optical patch cord.

### Fiber photometry experimental design

GCaMP7b was excited with a blue LED (Doric, LED driver, assembled with 470 nm) at 480 Hz, and emission was sampled at 2’000 Hz with a photodetector (Doric, DFD_FPA_FC) through a fluorescence MiniCube (Doric, ilFMC6_IE(400-410)_E1(460-490)_F1(500-540)_E2(555-570)_F2(580-680)_S) and digitized with a national instruments USB-6002 DAQ device. Fiber photometry recordings were acquired using custom-written scripts in LabVIEW on a computer. All the recordings were started about 15 min before ZT14 to stabilize the fluorescent signal. For every trial at ZT14 a constant light pulse of 10000 Lux (Daylight Lamp) was manually turned on for 15 min (± 20 seconds), and the recording was stopped 15 minutes after the light was switched off. To allow mice to restore their circadian time to the 12-hour light-dark cycle, the intertrial interval was at least 10 days. A patch cord was connected to the light source and the photometry system to align the light pulse to the photometry recording.

### In vivo calcium imaging, data processing and analysis

The data were subdivided into control (*scramble*) and experimental (*shCDK5*) groups. The fluorescent signal was demodulated in the frequency band of 470 – 490 Hz at 10 Hz acquisition rate. Due to GCaMP7b variable photobleaching (i.e., the loss of fluorescence intensity as a function of light exposure), we filtered the demodulated signal using a 3 order Savitzky-Golay filter (every 100 s), and detrended it using a hug-line. We then calculated the ΔF/F_0_ as follows:

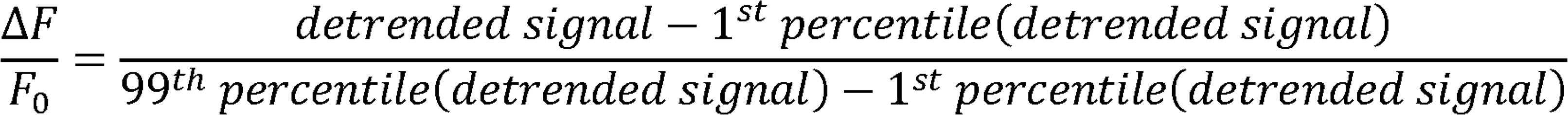

Finally, the ΔF/F_0_ was cut to the light pulse as follows: a) 5 minutes before the light pulse, b) 15 minutes during the light pulse and c) 15 minutes after the light. To exclude differences in the duration of the light pulse (± 20 seconds), the period analyzed was of 14 minutes. All data processing was performed using custom-written Matlab scripts.

### Live FRET imaging

The protocol was performed as before ^15^. The following plasmid was used for the project: ICAP-NLS Vector carrying ^15^. Transfected NIH3T3 cells were starved for 4 h with 0.5% FBS DMEM. Subsequently, cells were washed twice with 1×HBTS without CaCl2 and MgCl2 (25 mM HEPES, 119 mM NaCl, 6gr/L Glucose, 5 mM KCl) and resuspended in the same buffer. NIH3T3 cells were imaged using an inverted epifluorescence microscope (Leica DMI6000B) with an HCX PL Fluotar 5x/0.15 CORR dry objective, a Leica DFC360FX CCD camera (1.4 M pixels, 20 fps), and EL6000 Light Source, and equipped with fast filter wheels for FRET imaging. Excitation filters for CFP and FRET: 427 nm (BP 427/10). Emission filters for CFP: 472 (BP 472/30) and FRET: 542 nm (BP 542/27). Dichroic mirror: RCY 440/520. One frame every 20 sec was acquired for at least 90 cycles (0.05 Hz frequency), and the recording lasted at least 30 min. The baseline response in the presence of HBTS was recorded for 2 min and 40s. At minute 3:00, 100 µM Forskolin, 2 mM CaCl2, and 2 mM MgCl2 were added to the cells. The time-lapse recordings were analyzed using LAS X software (Leica). Two regions of interest (ROI) were randomly selected for each cell, and 50 cells per plate were chosen randomly. A first ROI delimiting the background and a second ROI including the cell nucleus of NIH3T3 cell expressing NLS KIDKIX were used per cell. The ROI background values were subtracted from the ROI cell values for each channel. For baseline normalization, the FRET ratio R was expressed as a ΔR/R, where ΔR is R–R0, and R0 is the average of R over the last 120 s prior stimulus.

### Statistical analysis

Statistical analysis of all experiments was performed using GraphPad Prism6 software. Depending on the data type, either an unpaired t-test or one- or two-way ANOVA with Bonferroni or Tukey’s post-hoc test was performed. Values considered significantly different are highlighted. [p<0.05 (*), p<0.01 (**), or p<0.001 (***)].

Data were compared via two-way repeated-measures ANOVA with post hoc Bonferroni’s corrections for multiple comparisons. Data distribution was assumed to be normal, but this was not formally tested. All data are displayed as mean ± standard error of the mean (SEM). No power calculations were performed to determine sample sizes, but similarly sized cohorts were used in previous studies. The experimenters were not blind to the conditions when acquiring or analyzing the data.

Sample numbers are indicated in the corresponding figure legends, and test details are only reported for significant results. Figures were prepared in Adobe Illustrator 2022.

## Acknowledgments

Technical assistance by Antoinette Hayoz and Maude Marmy is acknowledged. This work was supported by the Swiss National Science Foundation (SNF) 310030_219880/1 to UA, 310030_197607 to DAG, 310030_219438/1 to ZY, the Inselspital University Hospital Bern, the European Research Council CoG-725850 to AA and the States of Berne and Fribourg.

## Author contributions

Conceived and designed the experiments: AB, UA. Performed the experiments: AB, MB, GS, JR.

Analyzed the data: AB, MB, GS, JR, DG, AA, UA.

Contributed reagents, materials, analysis tools: DG, ZY, AA, UA Wrote the paper: AB, UA

## Competing interests

We declare no conflict of interest.

## Notes

### Competing Interest Statement

The authors have declared no competing interest.

### Summary of Updates

Figures 2, 4, 6, and 7 have been adapted Supplemental Figures Fig. 1, 2 contain additional data Supplemental Fig. 4 shows all the traces analysed The presentation of Fig. 7 shows original blots

