## Supplemental data for "Cyclin-dependent kinase 5 (Cdk5) activity is modulated by light and gates rapid phase shifts of the circadian clock"

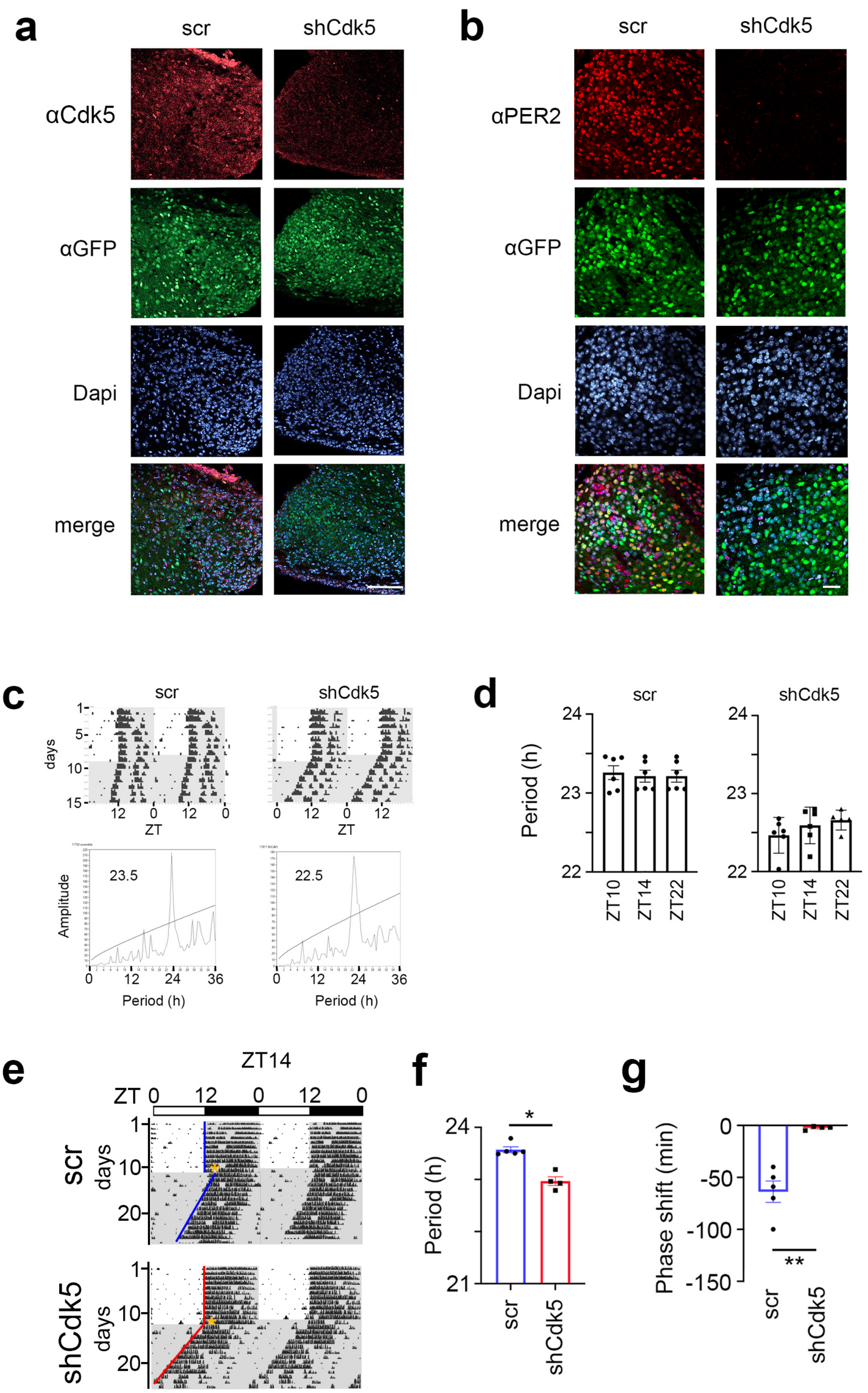


Supplementary Figure 1

**Supplementary Figure 1 Validation of knock-down of Cdk5 expression.**

(**a**) Representative sections of the SCN region after injection of AAVs carrying either scrambled (scr) or shCdk5 shRNA. Slices were stained with DAPI to stain nuclei (blue), and anti-GFP (green) and anti-Cdk5 (red) antibodies. GFP shows those cells infected by the virus. Cdk5 was efficiently down-regulated in the SCN by shCdk5 but not by scr shRNA (red), which was as efficiently delivered as shCdk5 (green). Scale bar: 100 µm (**b**) Same as (a) but stained with anti-PER2 (red). PER2 was efficiently down-regulated in the SCN by shCdk5 but not by scr shRNA, which was as efficiently delivered as shCdk5 (green). Scale bar: 30 µm (**c**) Examples of double plotted actograms of control (scr) and Cdk5 knock-down (shCdk5) animals (top panels). Mice were kept under a 12 h light / 12 h dark cycle for 8-10 days before they were released into constant darkness (DD). In DD period was determined using $\chi^2$-periodogram analysis (bottom panels). (**d**) The period was determined after a light pulse at ZT10, ZT14, or ZT22. $\tau$ control ZT10: 23.26 ± 0.09 h, ZT14: 23.21 ± 0.08 h, ZT22: 23.21± 0.08 h; $\tau$ shCdk5 ZT10: 22.47 ± 0.09 h, ZT14: 22.59 ± 0.1 h, ZT22: 22.66 ± 0.06 h. Values are the mean ± SEM (n = 6). The period was not significantly altered by the light pulse itself neither in control (scr) nor in shCdk5 animals. Brown-Forsythe ANOVA test, $p > 0.05$, n = 6.

(**e**) Examples of double-plotted wheel-running actograms of control (scr) and Cdk5 knock-down (shCdk5) female mice. Animals were kept under a 12 h light / 12 h dark cycle (white and grey areas, respectively) (LD). After 10-12 days they received a 15 min. light pulse at ZT14 (yellow stars). After the light pulse animals were released into constant darkness (DD). (**f**) The circadian period ($\tau$) of female shCdk5 mice (red) is significantly shorter compared to female scr controls (blue). $\tau$ scr = 23.53 ± 0.05 h, $\tau$ shCdk5 = 22.95 ± 0.08 h. All values are mean ± SEM, Mann Whitney test, n = 5 scr, n = 4-5, *$p < 0.02$. (**g**) Quantification of phase shifts ($\varphi$) after a 15 min. light pulse at ZT14. The phase shift at ZT14 is strongly reduced in shCdk5 animals (red) compared to scr controls (blue). scr: $\varphi$ ZT14: -63.8 ± 11.2 min., shCdk5: $\varphi$ ZT14: -2.5 ± 0.13 min. All values are mean ± SEM, unpaired t-test with Welch's correction, n = 4-5, **$p < 0.01$.

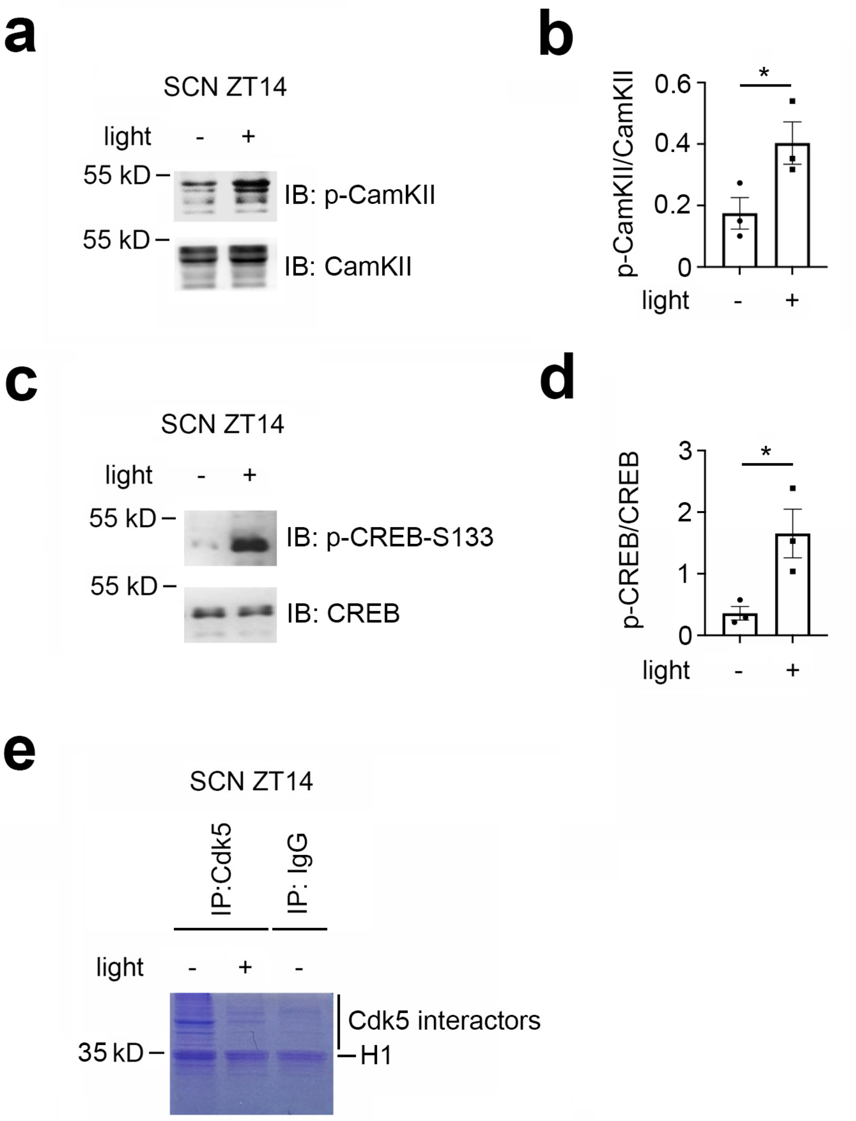


Supplementary Figure 2

**Supplementary Figure 2 Light-induced phosphorylation of CaMKII and control immunoprecipitation.** (**a**) Western blot of SCN tissue extracts at ZT14 with and without a light pulse. The lower panel shows the total CaMKII that was recognized by a polyclonal antibody. The upper panel shows increased levels of phosphorylated CaMKII (p-CaMKII) after light administration at ZT14. (**b**) Quantification of p-CaMKII relative to total CaMKII. Values are the mean ± SEM. Unpaired t-test, $n = 3$, $^*p < 0.05$. (**c**) Western blot of SCN tissue extracts at ZT14 with and without a light pulse. The lower panel shows the total CREB that was recognized by a polyclonal antibody. The upper panel shows increased levels of CREB phosphorylated at Ser-133 (p-CREB-S133) after light administration at ZT14. (**d**) Quantification of p-CREB-S133 relative to total CREB. Values are the mean ± SEM. Unpaired t-test, $n = 3$, $^*p < 0.05$. (**e**) SDS-Gel after immunoprecipitation of the SCN extracts with anti-Cdk5 and anti-IgG, respectively.

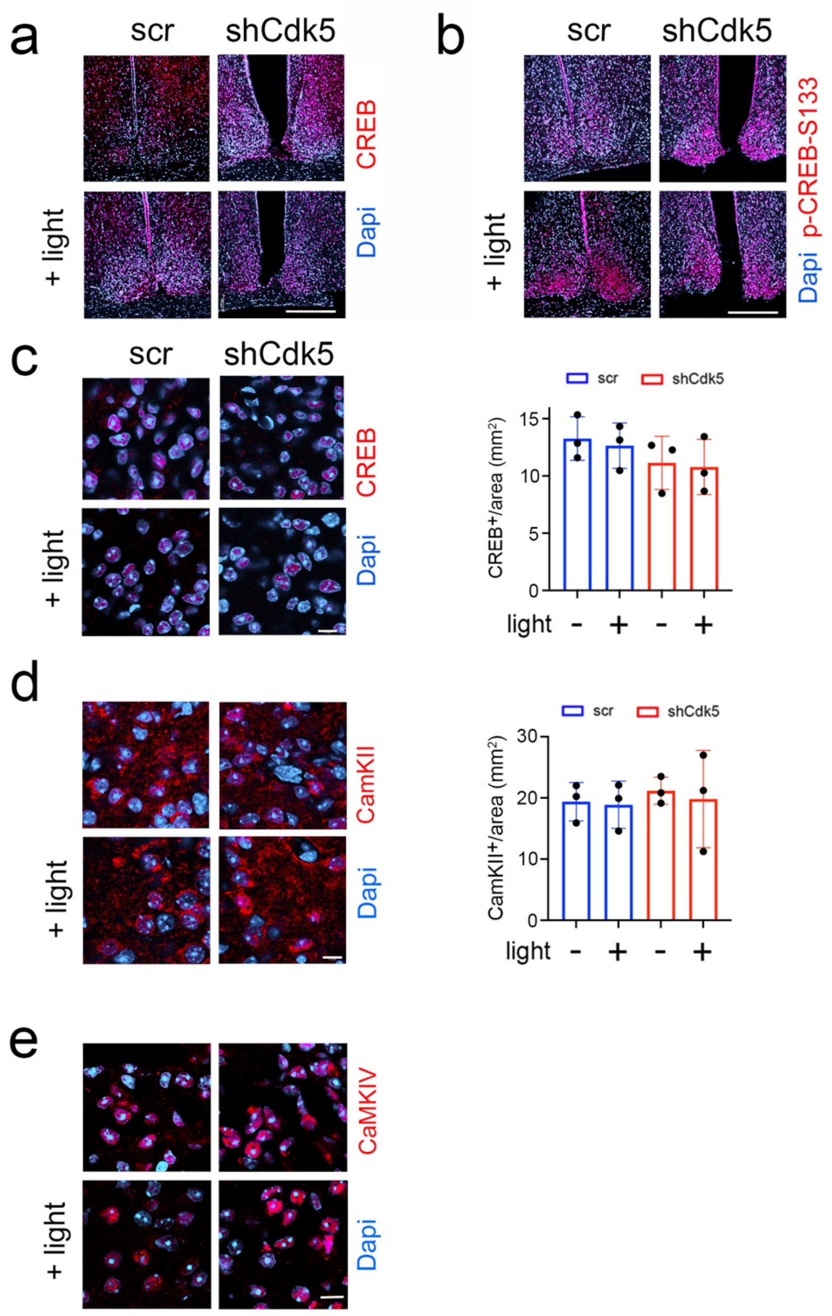


Supplementary Figure 3

**Supplementary Figure 3 Lower magnification images of SCN sections and quantification of total CREB, CaMKII, and CaMKIV.** (**a**) Immunohistochemistry on the SCN of control (scr) and shCdk5 mice using an antibody recognizing total CREB before and after a light pulse at ZT14. The red color shows CREB and the blue color represents Dapi-stained nuclei of SCN cells. Scale bar: 250 µm. (**b**) Immunohistochemistry on the SCN of control (scr) and shCdk5 mice using an antibody recognizing phospho-serine 133 of CREB (p-CREB-S133) before and after a light pulse at ZT14. The red color shows p-CREB-S133 and the blue color represents Dapi-stained nuclei of SCN cells. Scale bar: 250 µm. (**c**) Left panel: Same as (a) but at higher magnification. Right panel: Quantification of total CREB in scr (blue) and shCdk5 (red) SCN. Values are the mean ± SEM. Unpaired t-test, n = 3. Scale bar: 10 µm (**d**) Left panel: Immunohistochemistry on the SCN depicting total CaMKII (red) and cell nuclei (blue). Right panel: Quantification of total CaMKII in scr (blue) and shCdk5 (red) SCN. Values are the mean ± SEM. Unpaired t-test, n = 3. Scale bar: 10 µm. (**e**) Left panel: Immunohistochemistry on the SCN depicting total CaMKIV (red) and cell nuclei (blue). Right panel: Quantification of total CaMKIV in scr (blue) and shCdk5 (red) SCN. Values are the mean ± SEM. Unpaired t-test, n = 3. Scale bar: 10 µm.

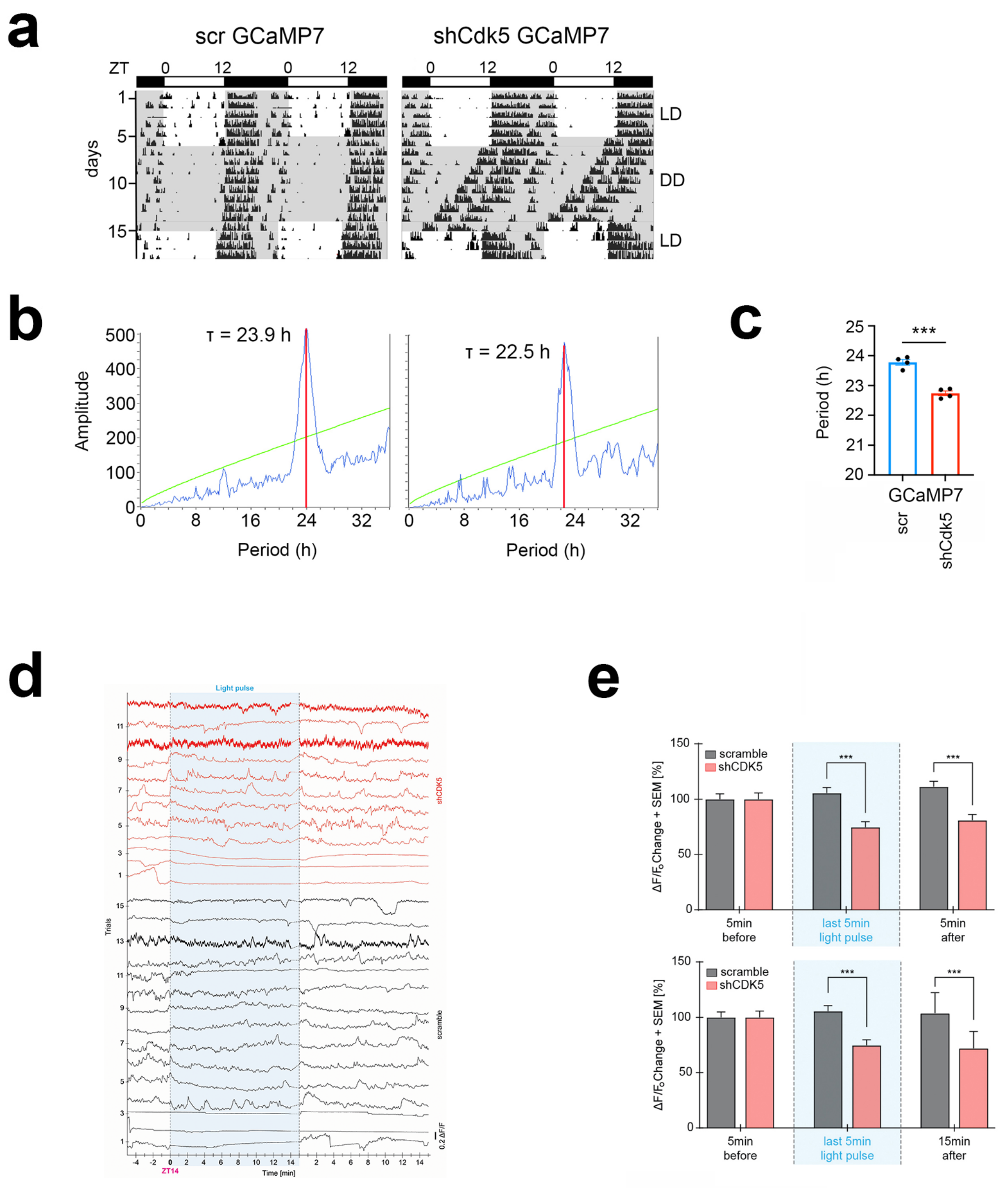


Supplementary Figure 4

**Supplementary Figure 4 GCaMP7 reporter does not affect the knock-down of Cdk5 in the SCN.** (**a**) Examples of double-plotted wheel-running actograms of control (scr) and Cdk5 knock-down (shCdk5) mice containing the reporter GCaMP7. Animals were kept under a 12 h light / 12 h dark cycle (white and grey areas, respectively) (LD). After 6 days they were released into constant darkness (DD). (**b**) $\chi^2$-periodogram analysis showing circadian period ($\tau$) for control GCaMP7 and shCdk5 GCaMP7 mice, τ scr = 23.9 h, τ shCdk5 = 22.5 h. (**c**) Quantification of circadian period (τ). τ scr = 23.78 ± 0.08 h, τ shCdk5 = 22.74 ± 0.09 h. All values are mean ± SEM, unpaired t-test with Welch's correction, n = 5-6, ***$p < 0.001$. (**d**) The activity of SCN neurons expressing GCamP7b (normalized $\Delta F/F$) in individual trials in the dark phase, 5 min. before and 15 min. after the 15 min. (± 20 s) light pulse delivered at ZT14. The 15th minute of the light recording has not been analyzed to homogenize the lightning time precisely. Black = scramble, N = 15 trials, n = 5 mice / red = shCdk5, N = 12 trials, n = 4 mice.

(**e**) Bar plots showing the percentage of $\Delta F/F_0$ changes ± SEM (normalized to the 5 minutes before the light pulse) in the dark phase, 5 minutes (top) or 15 minutes (bottom) after the light pulse delivered at ZT14 and in the last 5 minutes of the light pulse (min 10-14). Black: scramble, N = 15 trials, n = 5 mice; red: shCdk5, N = 12, trials, n = 4 mice. ***$p < 0.001$; Two-way ANOVA corrected with Bonferroni post-hoc test.

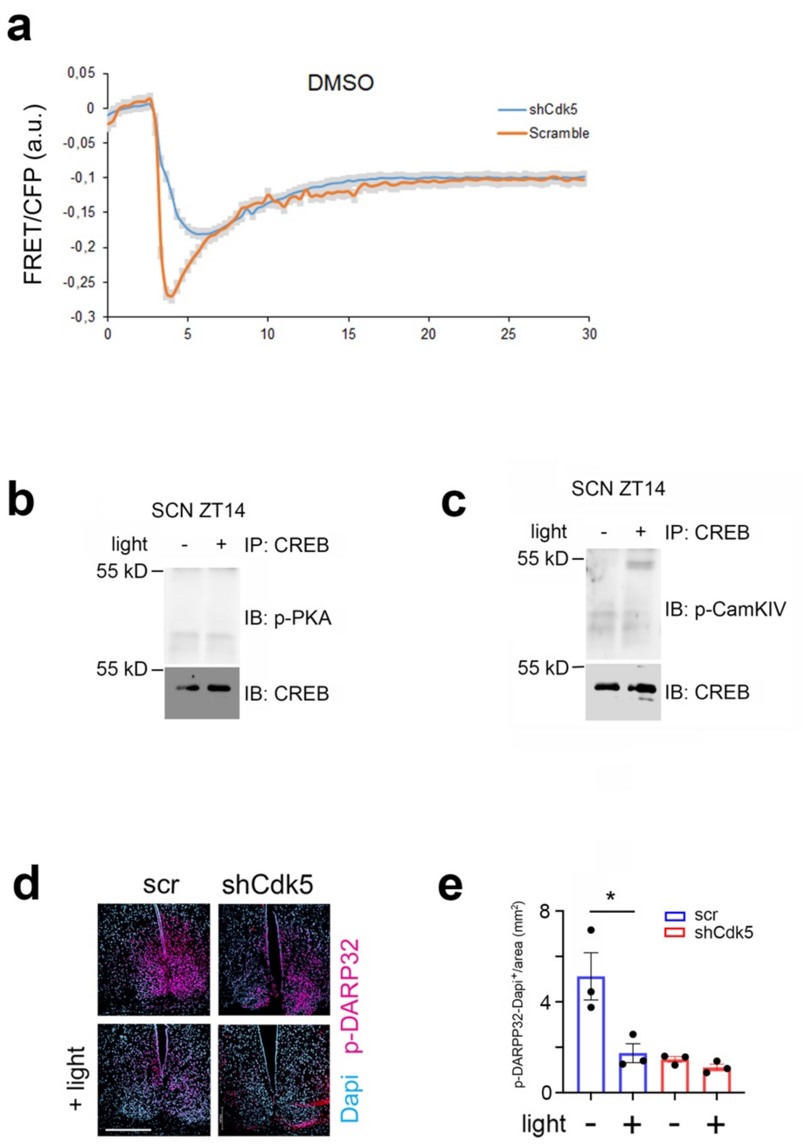


Supplementary Figure 5

**Supplementary Figure 5 Solvent control for FRET experiment. Immunoprecipitation of CREB and low magnification of SCN stained with p-DARPP32. (a)** FRET/CFP signal ratio changes with DMSO treatment in NIH 3T3 cells transfected with either a scr control (red) or shCdk5 (blue) expression construct. Values are the mean ± SD. Two-way ANOVA revealed no difference between the curves, n = 3. (**b**) Immunoprecipitation (IP) of CREB from SCN tissue at ZT14 before and after a light pulse. Immunoblot (IB) shows no interaction with p-PKA, IB: CREB loading control. (**c**) Immunoprecipitation (IP) of CREB from SCN tissue at ZT14 before and after a light pulse. Immunoblot (IB) shows interaction with p-CaMKIV, IB: CREB loading control. (**d**) Immunohistochemistry on the SCN of control (scr) and shCdk5 mice using an antibody recognizing p-DARPP32 before and after a light pulse at ZT14. The red color shows p-DARPP32 and the blue color represents Dapi-stained nuclei of SCN cells. Scale bar: 250 µm. (**e**) Quantification of nuclear p-DARPP32 signal. Values are the mean ± SEM. Unpaired t-test with Welch's correction, n = 3, *$p < 0.05$.

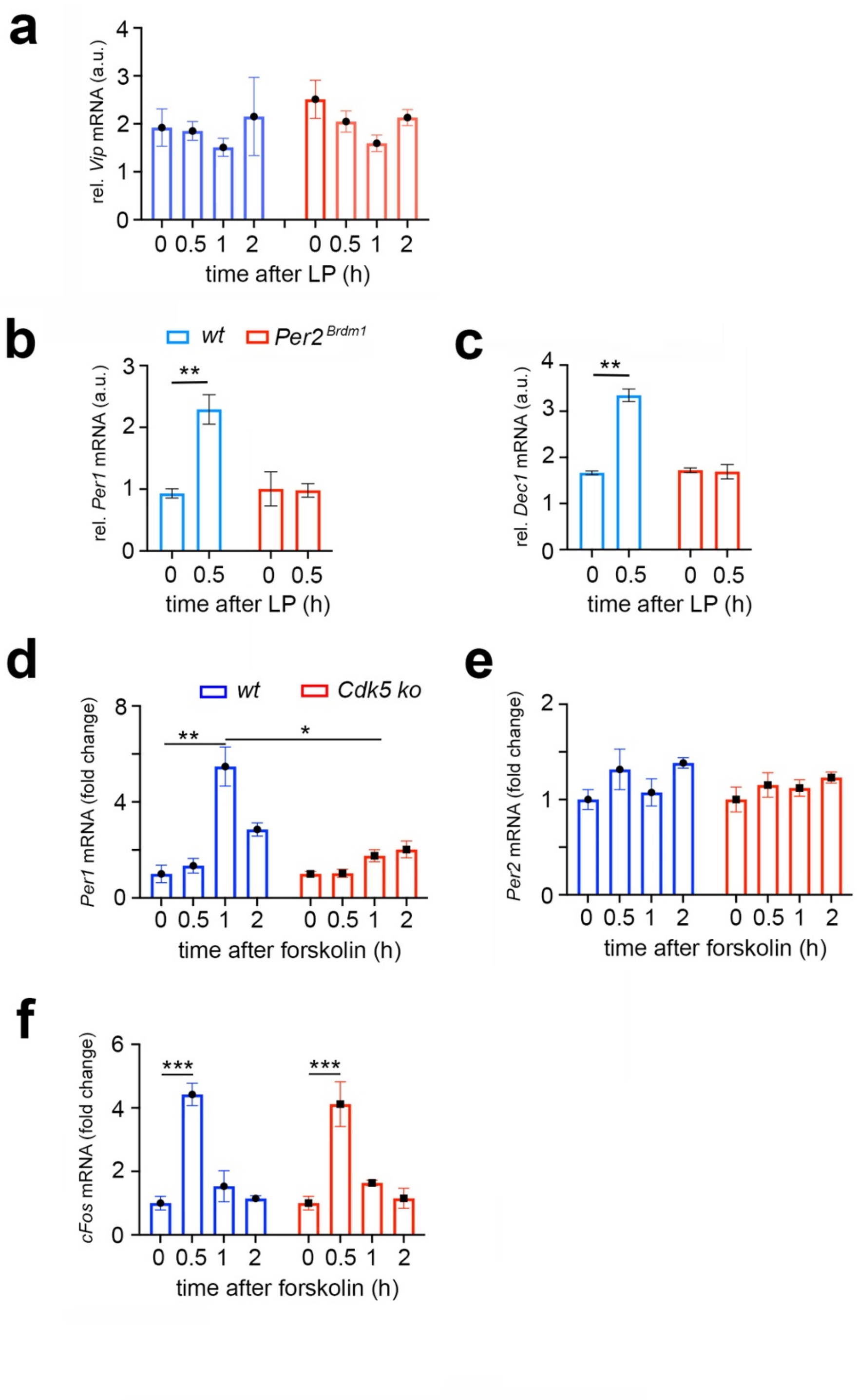


Supplementary Figure 6

**Supplementary Figure 6 *Vip*, *Per1, Dec1* levels in the SCN of shCdk5 and *Per2$^{Brdm1}$* mice and *Per1, 2* and *cFos* gene expression in NIH 3T3 cells. (a)** *Vip* mRNA expression is not induced by light in the SCN of scr control and shCdk5 animals. Scr: 0 h: 1.92 ± 0.39, 0.5 h: 1.85 ± 0.20, 1 h: 1.51 ± 0.19, 2 h: 2.15 ± 0.82; shCdk5: 0 h: 2.51 ± 0.40, 0.5 h: 2.05 ± 0.22, 1 h: 1.59 ± 0.17, 2 h: 2.13 ± 0.17. Values are the mean ± SEM. Unpaired t-test, n = 3. (**b**) Relative mRNA values are represented as blue bars for scr control animals and as red bars for *Per2$^{Brdm1}$* mice. The values were determined at 0 and 0.5 hours after a light pulse (LP) was given at ZT14. Induction of *Per1* mRNA expression by light 0.5 hours after light in wt control animals. In contrast, *Per1* is not induced in *Per2$^{Brdm1}$* SCN. wt: 0 h: 0.93 ± 0.07, 0.5 h: 2.29 ± 0.24; *Per2$^{Brdm1}$*: 0 h: 1.00 ± 0.28, 0.5 h: 0.98 ± 0.11. Values are the mean ± SEM. Unpaired t-test, n = 3, **p < 0.01. (**c**) Induction of *Dec1* mRNA expression 0.5 hours after light in wt control animals. In contrast, *Dec1* is not induced in the *Per2$^{Brdm1}$* SCN. wt: 0 h: 1.66 ± 0.04, 0.5 h: 3.35 ± 0.14; *Per2$^{Brdm1}$*: 0 h: 1.72 ± 0.05, 0.5 h: 1.69 ± 0.16. Values are the mean ± SEM. Unpaired t-test, n = 3, **p < 0.01. **(d)** Relative mRNA values are represented as blue bars for control wild-type (wt) cells and as red bars for Cdk5 knock-out (Cdk5 ko) cells. The values were determined 0, 0.5, 1, and 2 hours after forskolin treatment. Induction of *Per1* mRNA expression is maximal 1 h after forskolin treatment in wt cells, but not in Cdk5 ko cells. Wt: 0 h: 1 ± 0.36, 0.5 h: 1.34 ± 0.30, 1 h: 5.48 ± 0.81, 2 h: 2.85 ± 0.28; Cdk5 ko: 0 h: 1 ± 0.10, 0.5 h: 1.03 ± 0.17, 1 h: 1.76 ± 0.25, 2 h: 2.02 ± 0.35. Values are mean ± SD. Unpaired t-test, *p < 0.05, **p < 0.01, n = 3. (**e**) *Per2* mRNA expression is induced neither in wt cells nor in Cdk5 ko cells. Wt: 0 h: 1 ± 0.10, 0.5 h: 1.32 ± 0.21, 1 h: 1.07 ± 0.14, 2 h: 1.38 ± 0.05; Cdk5 ko: 0 h: 1 ± 0.13, 0.5 h: 1.15 ± 0.13, 1 h: 1.12 ± 0.09, 2 h: 1.23 ± 0.06. Values are mean ± SD. Unpaired t-test, n = 3. (**f**) Induction of *cFos* mRNA expression is maximal 0.5 h after forskolin treatment in wt cells, and in Cdk5 ko cells. Wt: 0 h: 1 ± 0.21, 0.5 h: 4.42 ± 0.35, 1 h: 1.53 ± 0.49, 2 h: 1.15 ± 0.08; Cdk5 ko: 0 h: 1 ± 0.21, 0.5 h: 4.12 ± 0.71, 1 h: 1.64 ± 0.09, 2 h: 1.15 ± 0.32. Values are mean ± SD. Unpaired t-test, n = 3.

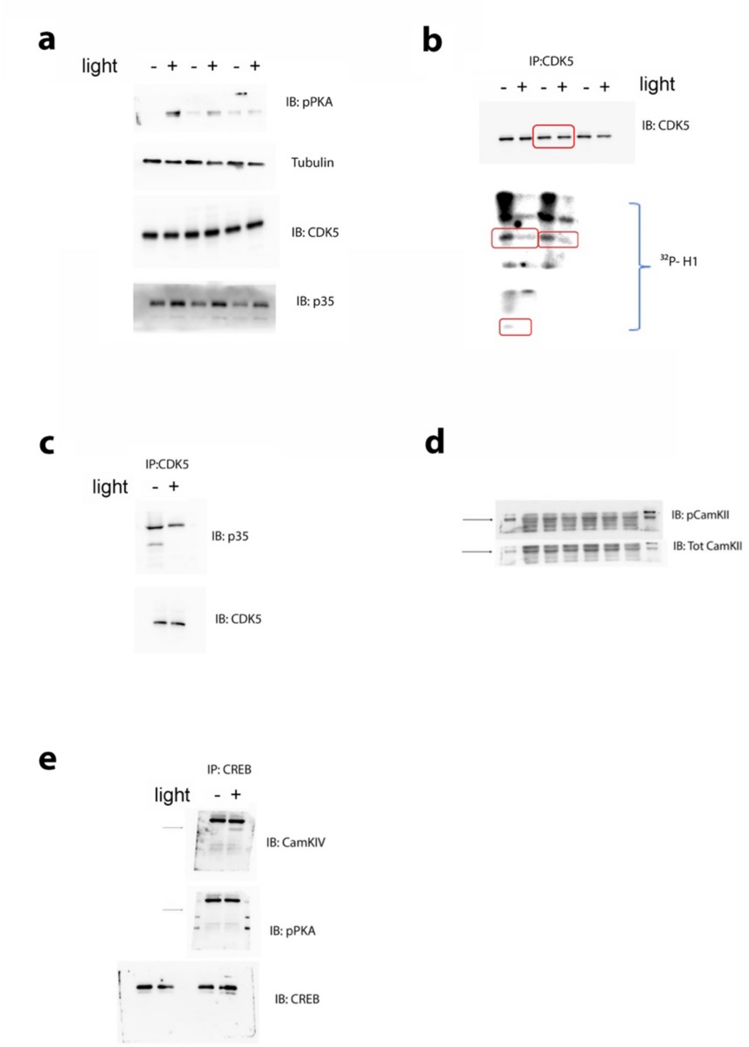


Supplementary Figure 7

**Supplementary Figure 7 Western blots used for quantification and full blots**
(**a**) Triplicates for quantification are shown in Figure 2c. (**b**) Triplicates for quantification are shown in Figure 2e. The red square in the top panel is the example shown in Figure 2d. Red squares in the bottom panel indicate the quantified H1 phosphorylation. (**c**) Full blot of Figure 2f (**d**) Triplicates for quantification shown in Supplementary Figure 2b (**e**) Full blots shown in Supplementary Figure 5b and 5c.
